# Strong selection at the level of codon usage bias: evidence against the Li-Bulmer model

**DOI:** 10.1101/106476

**Authors:** Heather E. Machado, David S. Lawrie, Dmitri A. Petrov

## Abstract

Codon usage bias (CUB), where certain codons are used more frequently than expected by chance, is a ubiquitous phenomenon and occurs across the tree of life. The dominant paradigm is that the proportion of preferred codons is set by weak selection. While experimental changes in codon usage have at times shown large phenotypic effects in contrast to this paradigm, genome-wide population genetic estimates have supported the weak selection model. Here we use deep genomic sequencing of two *Drosophila melanogaster* populations to measure selection on synonymous sites in a way that allowed us to estimate the prevalence of both weak and strong selection. We find that selection in favor of preferred codons ranges from weak (|*N_e_s*| ∼ 1) to strong (|*N_e_s*| > 10). While previous studies indicated that selection at synonymous sites could be strong, this is the first study to detect and quantify strong selection specifically at the level of CUB. We suggest that the level of CUB in the genome is determined by the proportion of synonymous sites under no, weak, and strong selection. This model challenges the standard Li-Bulmer model and explains some of the longest-standing puzzles in the field.

## 2 Introduction

The degeneracy of the genetic code leads to protein-coding mutations that do not affect amino acid composition. Despite this, such synonymous mutations often have consequences on phenotype and fitness. The first evidence of the functionality of synonymous sites came from the discovery of codon usage bias (CUB), where, for a given amino acid, certain codons are used more frequently in a genome than expected by chance (Ikemura 1981; Grantham *et al.* 1981). The consensus in the field is that CUB is often driven by natural selection but the nature and strength of natural selection acting to maintain CUB is disputed.

The most common explanations for CUB postulate selection on either the rate or the accuracy with which ribosomes translate mRNA to protein. The existence of selection at synonymous sites at the level of translation is supported by several key observations. First, the preference toward particular “preferred” codons is consistent across genes within a particular genome suggesting a global, genome-wide process and not preference for the use of particular codons within specific genes (Chen 2004; Grantham 1980). Second, optimal codons tend to correspond to more abundant tRNAs, suggesting a functional relationship between translation and CUB (Post *et al.* 1979; Ikemura 1981; Ikemura 1982; Qian 2012). Third, preferred codons are more abundant in highly expressed genes than in the rest of the genome (Gouy 1982; Bulmer 1991; Novoa & Ribas de Pouplana 2012), consistent with selection being proportional to mRNA transcript abundance. Finally, constrained amino acid positions tend to contain preferred codons more frequently, suggesting a link between CUB and translational accuracy (*Escherichia coli*: Stoletzki & Eyre-Walker 2007; *Drosophila melanogaster*: Akashi 1994; mammals: Drummond & Wilke 2008). In addition to speed and accuracy, there is evidence that other processes are affected by codon composition, such as cotranslational folding (Pechmann & Frydman 2013), RNA stability (Presnyak *et al.* 2015), and transcription (Carlini & Stephan 2003; Newman *et al.* 2016; Zhou *et al.* 2016).

Beyond the level at which selection operates to generate CUB, it is important to consider how strong selection at synonymous sites is likely to be. This question has been most thoroughly addressed with population genetics approaches introduced by seminal papers of Li and Bulmer (Bulmer 1991; Li 1987). The Li-Bulmer model proposes that the observed proportion of preferred codons can be explained by the balance of mutation, selection (in favor of preferred codons), and random genetic drift. This model assumes a constant selection coefficient per codon or codon preference group. Given that levels of CUB are intermediate, even for highly biased genes of species with pronounced CUB, the model predicts that the strength of selection in favor of preferred codons should be on the order of the reciprocal of the effective population size (*N_e_s* ∼ −1). Indeed, if selection was an order of magnitude stronger, we expect > 99% of synonymous sites to be fixed at the preferred state. If it was an order of magnitude weaker, we would see no CUB at all (Hershberg & Petrov 2009).

The predicted weak selection should be detectable as a slight deviation in the site frequency spectrum (SFS). Mutations from preferred to unpreferred codons should reach comparatively lower frequencies in the population than those in the opposite direction. Such deviations have in fact been observed in many organisms that show clear CUB (*D. melanogaster*: Zeng *et al.* 2009; *Caenorhabditis remanei*: Cutter & Charlesworth 2006; *E. coli*: Sharp *et al.* 2010). These findings solidified the conclusion that selection at synonymous sites is weak but detectable.

Li-Bulmer model further predicts that the level of CUB should be very sensitive to the variation in effective population size. Such variation can be driven either by demographic differences between species or various levels of linked selection within genomes. There is indeed some equivocal evidence that species with higher effective population sizes do exhibit higher CUB. For instance, the CUB is stronger in *D. simulans* compared to *D. melanogaster*, which has a smaller effective population size (Akashi 1996, Jackson *et al.* 2017). There is also stronger CUB in *Drosophila* (large *N*_*e*_) than in mammals (small *N*_*e*_) (Urrutia 2001). However, there appears to be no correlation between levels of effective population size across mammals and levels of CUB (Kessler 2014). And neither does CUB appear to consistently correlate with levels of genetic draft within genomes. For instance, there is no consistent relationship between recombination rate and CUB (Singh 2005; Campos *et al.* 2013), and the correlations that have been identified (Kliman & Hey 1993; Campos *et al.* 2012) can alternatively be explained by variation in mutational biases (Marais 2001). Most importantly, the observed variation in the levels of CUB across the genome is minor at best, whereas the Li-Bulmer model of constant weak selection in favor of preferred codons predicts that codon bias should be exponentially sensitive to *N*_*e*_. Thus, the levels of CUB should be varying from nonexistent in the areas of no recombination to complete in the areas of high recombination.

An additional reason for the popularity of the Li-Bulmer estimate of weak selection driving CUB is that it matches the intuition that a synonymous change should not have a large phenotypic effect. However, there is abundant experimental evidence that this is not always the case. For example, optimizing the codon composition of the viral protein BPV1 increases the heterologous translation of the protein in humans by more than 1000 fold (Zhou *et al.* 1999). In *D. melanogaster*, changing a small number of preferred codons to unpreferred codons in the alcohol dehydrogenase (*Adh*) gene results in substantial changes in gene expression and in ethanol tolerance (Carlini & Stephan 2003, 2004). These studies suggest that some synonymous codon changes are subject to an *N*_*e*_*s* >> 1 (*s* >> 10^−6^). Similarly, mechanistic models of codon usage bias suggest that selection at preferred sites should often be several orders of magnitude larger than predicted by the population genetic Li-Bulmer model. In fact, the original Bulmer 1991 paper made this point and presented the discrepancy as a puzzle to be solved.

How can we reconcile these various lines of evidence? One possibility is that different synonymous codons are subject to very different levels of selection, with some subject to very strong selection and thus fixed at preferred states, some being neutral and found in mutational equilibrium, and some subject to weak selection and giving observed deviations in the SFS. This possibility would explain how the level of CUB can be intermediate in many genes, show deviations in the SFS consistent with weak selection, and at the same time be insensitive to deviations in effective population size.

This model can be tested using population genetic data. However, rather than consider only the shape of the SFS at synonymous sites in shallow population samples, which is what is usually done, we must either measure it in extremely deep samples or assess the overall level of polymorphism at synonymous sites as well. More precisely, one commonly used method of estimating level of selection is to compare the shape of the SFS at a putatively selected class of sites to that of a neutral reference. This approach is powerful, as the neutral reference can make the test independent of the demographic history of a population. Such tests have been used to estimate the strength of selection on codon bias in *D. melanogaster* and have failed to find any evidence of strong selection on codon bias (Clemente & Vogl 2012a; Singh *et al.* 2007; Zeng & Charlesworth 2009; 2010, Campos *et al.* 2013). However, the limit of detection in the aforementioned studies was set by the lowest allele frequency class in the dataset (set by the number of individuals sampled). As strong purifying selection results in a enrichment of very low allele frequency variants, only very deep population sequencing would allow for the detection of strong purifying selection. In the absence of very deep and accurate population sequencing, an alternative method is to utilize information about the proportion of sites that are polymorphic (polymorphism-level). Since both strong purifying selection and a decreased mutation rate can lower the polymorphism-level, the selected class of sites would have to be compared with a neutral reference that is matched for mutation rate and levels of linked selection. If the proportion of sites under strong selection is not very large, such approaches also require much larger amounts of genomic data than was available previously.

Intriguingly, a study by Lawrie *et al.* (2013) that did incorporate polymorphism-level and SFS with the use of matched neutral controls did find evidence of strong purifying selection. However, Lawrie and colleagues focused on selection on synonymous sites in general and failed to detect substantial selection on CUB. The Lawrie *et al.* 2013 study may have been limited in power due to the depth of population sequencing, a lack of ancestral polarization, a focus on highly conserved genes, and the use of a bottlenecked population (resulting in fewer variants).

Here we test for strong purifying selection on CUB in two distinct *D. melanogaster* populations. We accomplish this by comparing the polymorphism-level and SFS of fourfold degenerate synonymous sites in preferred and unpreferred codons to that of a short intron neutral reference. The neutral reference is produced by matching each fourfold site to a short intron site that is located within 1kb and has the same nucleotide at the position of interest and at the 5’ and 3’ neighboring sites. This creates a neutral reference that is subject to the same mutation rate and environment of linked selection as the fourfold sites. We find evidence that the there is a distribution of selection strengths on CUB, ranging from weak to strong. Our findings of strong selection on CUB directly conflict with previous models of codon bias that predict uniformly weak selection and indicate that the functional effects of CUB have been generally underestimated.

## 3 Results

### 3.1 Sequence data and neutral controls

We identified all fourfold degenerate synonymous (4D) sites and putatively neutral short intron (SI) sites in two datasets, one of an African (Zambia) and one of a North American (DGRP Freeze 2) *D. melanogaster* population. Each dataset consisted of ∼200 individual full-genome sequences. We down-sampled to 160 individuals per site per population to ensure equivalent power at each site. In order to reduce the effect of sequencing error, we filtered out low-quality bases (MAPQ < 20) for each individual genome sequence. We also excluded sites within 10bp of an indel. Since mapping errors are more common in regions around indels, and since introns have a greater number of indels, including these regions would have artificially inflated the SI polymorphism level and would have resulted in overestimates of purifying selection in 4D sites.

We used short introns as our neutral reference, as *D. melanogaster* short introns have been previously found to be under minimal selective constraint (Haddrill *et al.* 2005; Parsch *et al.* 2010, Clemente & Vogl 2012). Specifically, we used short introns less than 86bp in length, excluding the first 16bp and the last 6bp of each intron (Halligan & Keightley 2006). We matched 4D sites to SI sites based on ancestral nucleotide (polymorphisms polarized using the *D. simulans* genome), mutational context (the same two flanking nucleotides), and location (within 1000bp). All sites without an appropriate match were discarded. As the number of 4D sites was greater than the number of SI sites, we allowed SI sites to be matched to multiple 4D sites. This resulted in a total of 1075K 4D sites matched to 319K SI sites for Zambia, and 1183K 4D sites matched to 378K SI sites for DGRP. We performed the 4D/SI matching 200 separate times, producing 200 SI control sets.

### 3.2 Some synonymous sites are under strong selection

In order to detect the presence of purifying selection on synonymous sites we compared the synonymous 4D site frequency spectrum (SFS) and polymorphism levels to that of the matched SI controls. Purifying selection removes genetic variation from a population, resulting in a decrease in the polymorphism-level (the proportion of polymorphic sites). The effect of purifying selection on the shape of the SFS is a function of the strength of selection. Weak purifying selection (*N*_*e*_*s* > −1) decreases the density of the SFS at intermediate allele frequencies and enriches low frequency variants. Strong purifying selection (*N*_*e*_*s* < −10) results in an enrichment of very low allele frequency variants, making a skew in the SFS detectable only when a large number of individuals have been sampled. The most extreme example of this is of lethal mutations (*N_e_s* = −*Inf*), which do not affect the shape of the SFS and result exclusively in a decrease in the polymorphism-level.

We first performed a maximum likelihood (ML) estimation of the strength and amount of selection on synonymous sites using only polymorphic sites. We call this the “shape-only” ML model because it relies solely on deviations in the shape of the SFS and does not use polymorphism-level information. In this analysis we tested both the full datasets and the subset of synonymous sites found in preferred codons (the most frequent codon per amino acid). We hypothesized that preferred codons were under stronger purifying selection than unpreferred codons. For both the Zambia and the DGRP full datasets, we found no evidence for selection. We performed the same test on the subset of 4D sites in preferred codons and found strong evidence for selection in the Zambia population, estimating that 28% (95% bootstrap CI: 26-29) of sites were under purifying selection at *N_e_s* = −3 (95% bootstrap CI: 1-6). For the DGRP population, the selection estimates were similar (20% at *N_e_s* = −4); however, the confidence intervals were much larger (18% − 94% at *N_e_s* = −*Inf* − 0) and the selection model was not significantly better than neutrality. One factor potentially contributing to the large confidence intervals for DGRP is reduced power due to fewer polymorphic sites (less than one-half the number of polymorphisms as Zambia). The detection of purifying selection in the set of 4D sites in preferred codons suggested that 1) synonymous sites in preferred codons were under greater purifying selection than the genomic average and 2) we were underpowered to detect this selection in the full dataset due to a low total proportion of sites under selection.

Strong purifying selection will result in an enrichment of rare alleles and an overall reduction in polymorphism. The amount of reduction in polymorphism in the 4D sites compared with the SI controls can be expressed as the “polymorphism ratio”, defined as the natural logarithm of the SI polymorphism to 4D polymorphism ratio. The polymorphism ratio is positive for a depletion of 4D polymorphism and negative for an excess of 4D polymorphism. We found a reduction in 4D polymorphism in both the Zambia and DGRP datasets, with a polymorphism ratio of 0.10 and 0.14, respectively (Figure 1). We found an even greater reduction in polymorphism in preferred codons, with a polymorphism ratio of 0.19 and 0.29 for Zambia and DGRP, respectively. The expected polymorphism ratio for Zambia preferred codons, based on the shape-only ML estimate of 28% sites under selection at *N_e_s* = −3, is 0.07 (95% CI estimate: 0.03-0.11). This expected polymorphism ratio of 0.07 was significantly lower than the observed polymorphism ratio of 0.19, suggesting that the shape-only ML model does not fully explain the data.

The strong reduction in 4D polymorphism is suggestive of strong selection operating on 4D sites. In order to measure strong selection on 4D sites we included the polymorphism-level in the ML selection estimate. We call this the “level + shape” ML model. We tested five different level + shape ML models: 1) neutral, 2) neutral + lethal, 3) neutral + 1 selection coefficient, 4) neutral + selection + lethal, and 5) neutral + 2 selection coefficients (Table 1; see Methods). For the Zambia dataset, the best fit model was the neutral + 1 selection coefficient model (12% at *N_e_s* = −20). For the set of preferred codons the best-fit model was the neutral + 2 selection coefficient model (16% at *N_e_s* = −23 and 44% at *N_e_s* = −1), indicating that there was a range of detectable selection coefficients acting at the preferred sites (Figure 1). The lack of a weak-selection estimate for the full dataset is consistent with the previous finding that the proportion of sites under weak selection is too low for detection when including all sites. For the full DGRP dataset, our level + shape ML selection estimate also detected strong selection (13% of sites at *N_e_s* = −86); however, the neutral + 1 selection coefficient model was not significantly better than the neutral + lethal model (13% lethal).

One factor contributing to the failure to significantly differentiate the strong selection class from lethality is reduced power due to fewer polymorphisms. For the number of polymorphisms in the DGRP dataset (∼ 47K), the strongest selection that should be distinguishable from lethality is *N_e_s* ∼ −60 (for power analyses, see Supplementary Figure 2). It is not surprising then that the selection model, with an *N_e_s* estimate of −86, was not significantly better than the lethal model. For unpreferred 4D sites, no selection model was significantly better than the neutral model, indicating a very low proportion of sites under selection. This is consistent with the low polymorphism ratios for these datasets (0.01 for both Zambia and DGRP). The enrichment of sites under selection in the set of preferred codons and the lack of selection found in the set of unpreferred codons indicates that selection on CUB is a major component of the total amount of purifying selection on synonymous sites, and that the identification of both a weak and a strong selection class for preferred codons indicates that selection on CUB may not be limited to weak selection, as generally believed.

**Figure 1:**
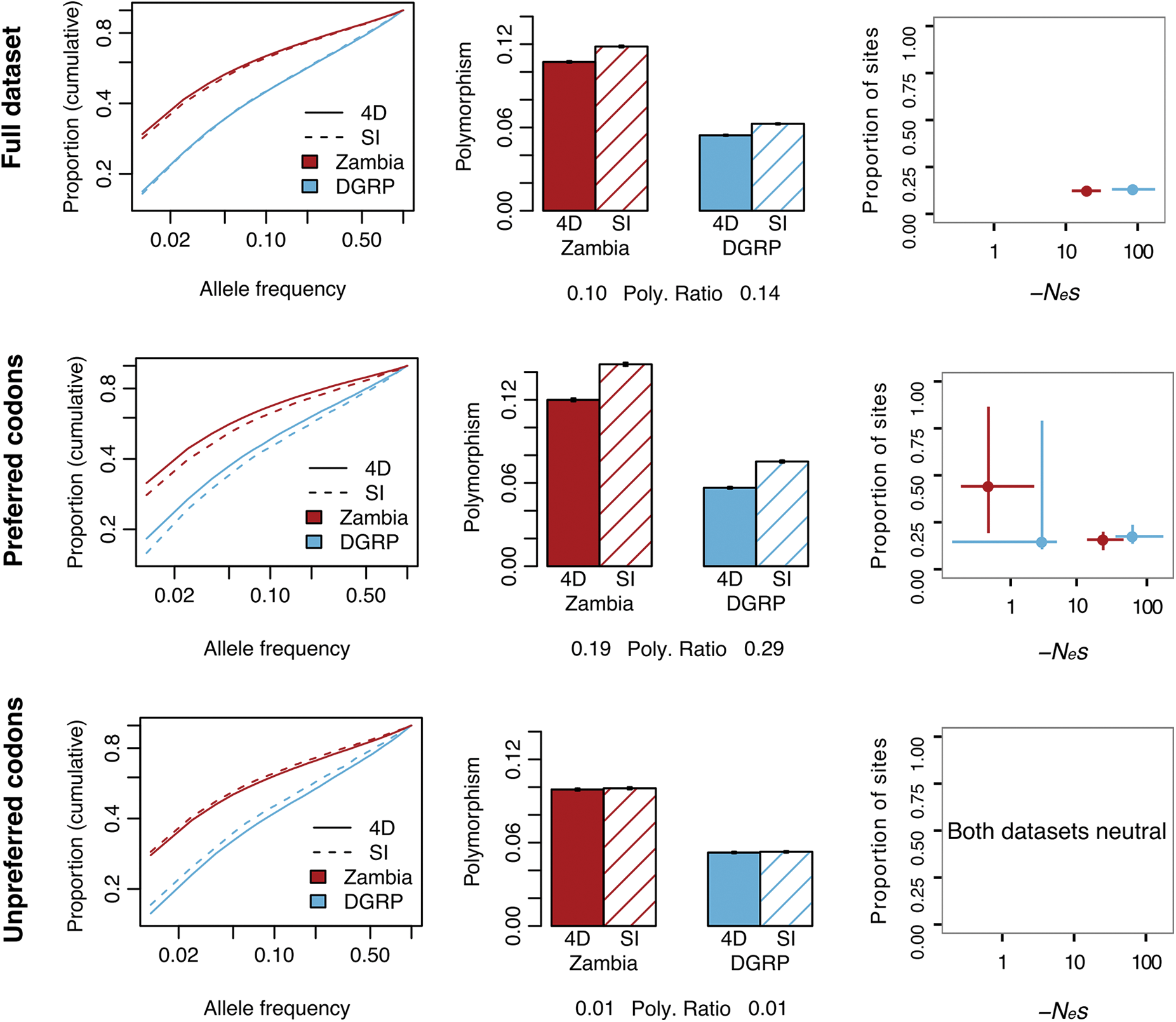
SFS (cumulative), polymorphism, and “level + shape” ML selection estimates for fourfold synonymous (4D) and matched short intron control (SI) sites for the full dataset (top), for preferred codons (middle), and for unpreferred codons (bottom).

**Table 1:**
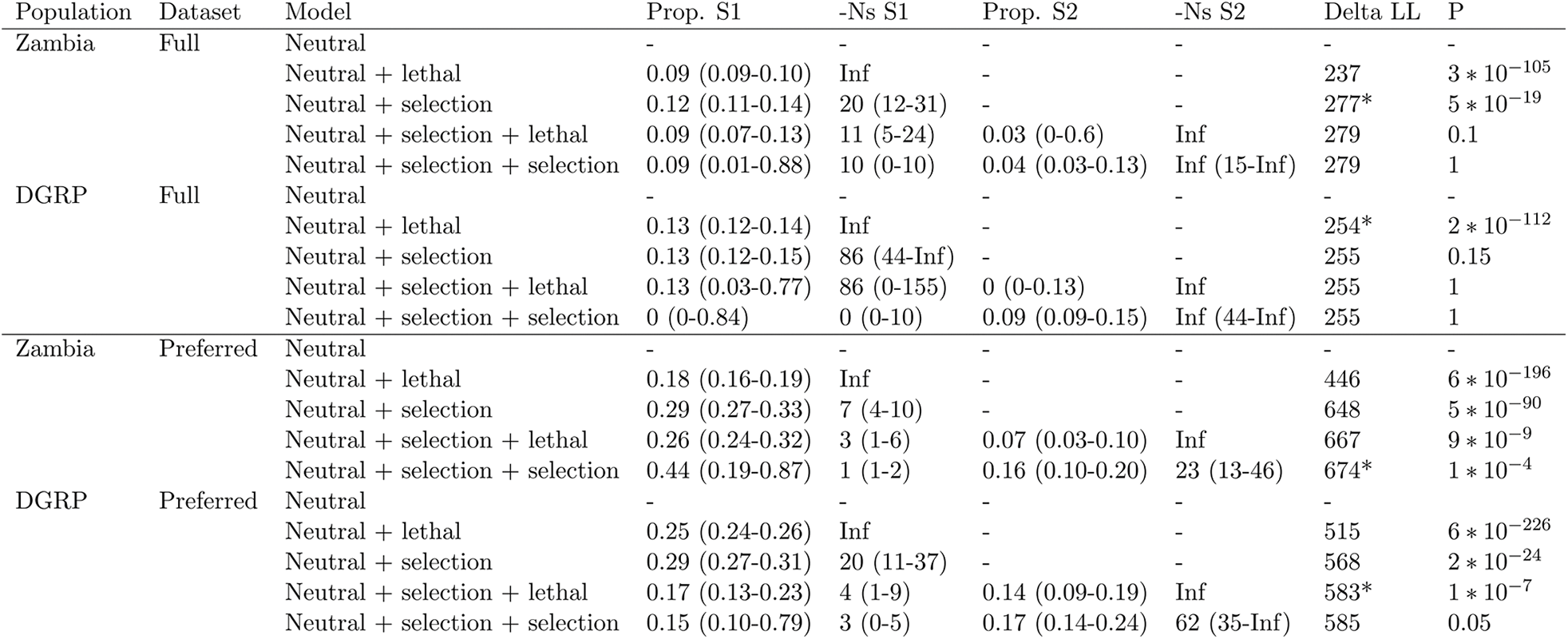
Nested “level + shape” maximum likelihood models tested for the Zambia and DGRP datasets. Values in parentheses are 95% bootstrap confidence intervals. Model comparison was performed with *chi*^2^ goodness of fit test *P* < 0.05: *Best fit model.

### 3.3 Phylogenetic conservation scores support finding of strong selection on CUB

If our ML and polymorphism ratio estimates truly do reflect selection levels, we might also expect our estimates to correlate well with signatures of long-term selection, such as phylogenetic conservation. We calculated phylogenetic conservation across a 10-species *Drosophila* phylogeny as the phyloP score from the program PHAST (Cooper *et al.* 2005). The phyloP conservation score measures the extent of conservation or divergence per site, with positive values representing conservation and negative values representing divergence. We excluded *D. melanogaster* from the phylogenetic analysis in order to avoid a confounding effect of *D. melanogaster* polymorphism on both polymorphism ratio and phyloP score. We asked if there was a correlation between the proportion of sites we identified to be under purifying selection and the level of phylogenetic conservation. We found a strong correlation between polymorphism ratio and phyloP conservation score of 4D sites (Zambia: *R*^2^ = 0.96, *P* < 2 ∗ 10^−16^; DGRP: *R*^2^ = 0.94, *P* < 2 ∗ 10^−16^) (Figure 2). We also performed level + shape ML estimates of the proportion of sites under selection for 4D sites in low (lower quartile), medium (middle two quartiles), or high (upper quartile) phyloP scores. Not only did we observe the same relationship of increasing purifying selection with increasing conservation, we also found that there was a tight correlation between the ML estimates of the proportion of sites under purifying selection and the polymorphism ratio (Supplemental Figure 3). The agreement of the polymorphism ratio and the level + shape ML estimates supports the use of polymorphism ratio as a rough proxy for the proportion of sites under strong purifying selection. The correlation of phylogenetic conservation with our estimates of purifying selection supports the relevance of our estimates to long-term constraint.

**Figure 2:**
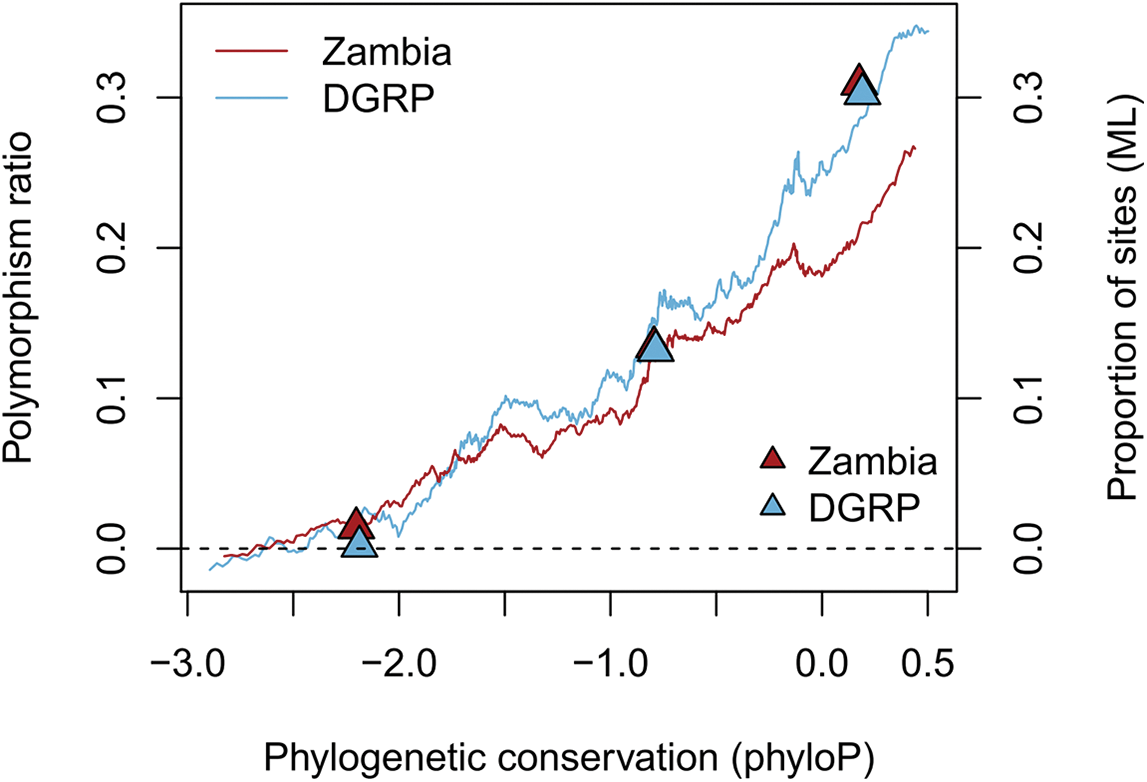
Correlation between the phyloP conservation score across a *Drosophila* phylogeny (excluding *D. melanogaster*) and the proportion of sites under selection, as estimated by the polymorphism ratio (lines) and the level + shape ML method (triangles). Dark red: Zambia, light blue: DGRP. The polymorphism ratio was estimated in sliding windows of 100K SNPs. The ML estimates were made for three groups: the lowest quartile, the middle two quartiles, and the highest quartile of phyloP scores. ML estimates are plotted against the median phyloP score for each group.

### 3.4 Recombination rate does not influence CUB

Previous studies have found evidence of only weak correlation between recombination rate and CUB. We tested for increased levels of purifying selection on preferred 4D sites as a function of recombination rate. We found that there was a greater proportion of preferred codons in high recombination rate regions (42.1% and 42.6% for Zambia and DGRP, respectively) than in low recombination rate regions (40.1% and 41.2% for Zambia and DGRP, respectively; both *chi*^2^ *P* < 10^−15^). However, once we controlled for mutational rates by measuring the polymorphism ratio, we found no evidence of increased strong purifying selection (greater polymorphism ratio) on preferred codons in high recombination rate regions compared with those in low recombination rate regions (Figure 3), nor any general increase in selection with recombination rate (Supplementary Figure 5).

**Figure 3:**
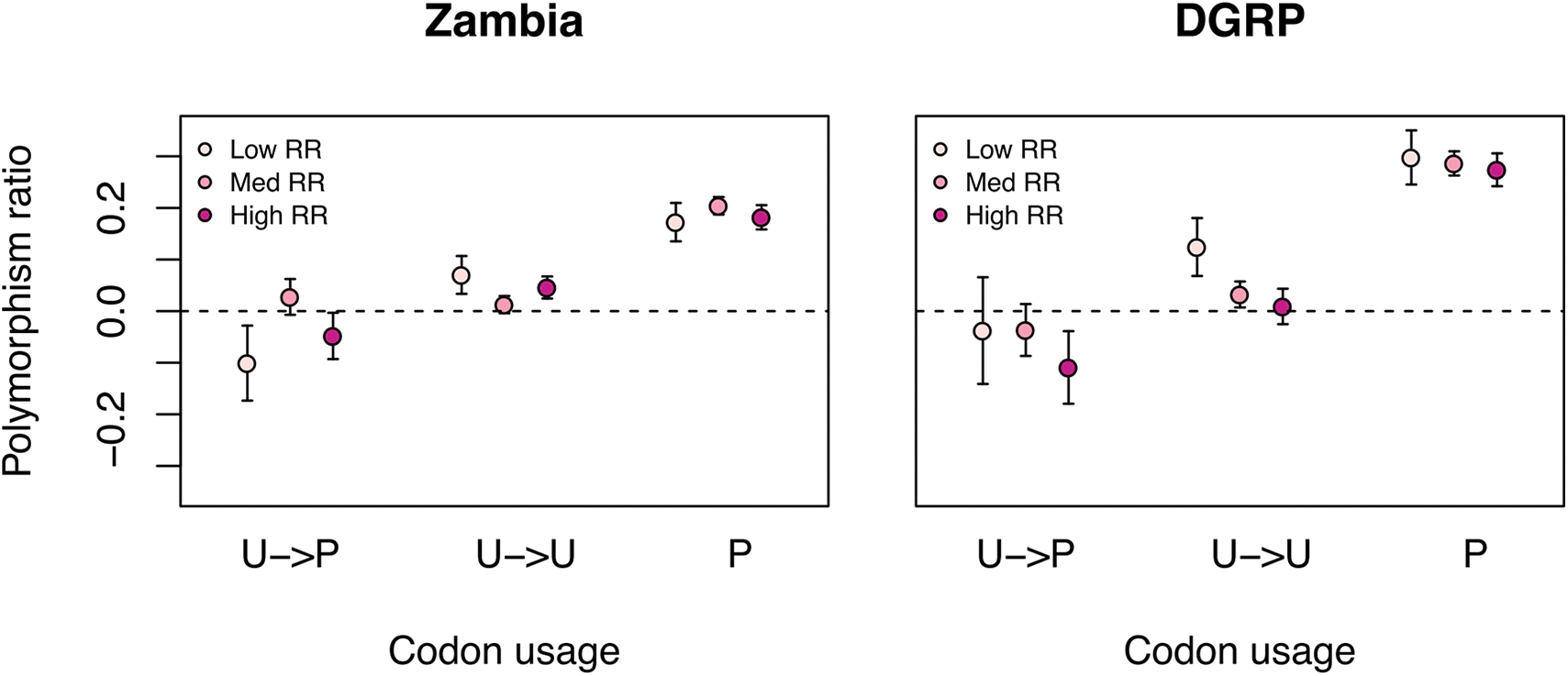
Polymorphism ratio by class of codon preference and recombination rate (RR). RR groups are classified into low (lowest quartile), medium (middle two quartiles), and high (top quartile). U: unpreferred; P: preferred. Error bars are 2 standard error.

### 3.5 Level of preference for a codon predicts proportion of sites under strong selection

Our findings suggested that a substantial proportion of synonymous sites in preferred codons were under strong purifying selection. Since the biased usage of codons actually exists on a continuum, rather than binary designations of “preferred” and “unpreferred”, we next asked whether or not the *level* of biased usage (for a particular codon) correlates with the amount of strong selection observed. We used the relative synonymous codon usage (RSCU) as a measure of the level of codon preference (Sharp & Li 1986). We measured RSCU for each 4D codon and compared that to the polymorphism ratio, which we take as a measure of the proportion of sites under strong selection. We found a strong positive relationship between RSCU and polymorphism ratio (Zambia: *R*^2^ = 0.56, *P* = 4∗10^−7^; DGRP: *R*^2^ = 0.63, *P* = 4∗10^−8^; Figure 4). We next asked if the change in RSCU, from ancestral to derived, correlated with polymorphism ratio. We hypothesized that mutations to a less preferred state (positive RSCU change) would show evidence for strong purifying selection (positive polymorphism ratio), whereas mutations to a more preferred state (negative RSCU change) would be positively selected for and have an increased level of 4D polymorphism relative to the SI control (negative polymorphism ratio). We found a strong, positive relationship between RSCU change and polymorphism ratio, with negative polymorphism ratios for strongly preferred derived mutations on unpreferred ancestral codons (Figure 4). This supports the hypothesis of purifying selection on the strongest unpreferred changes and positive selection on the strongest preferred changes. Note that negative polymorphism ratios (that is greater levels of polymorphism at 4D than SI sites), assuming that SI sites are neutral and 4D sites are under selection, is possible depending on the particulars of the mutational biases and direction of selection (Lawrie *et al.* 2011, McVean & Charlesworth 1999).

**Figure 4:**
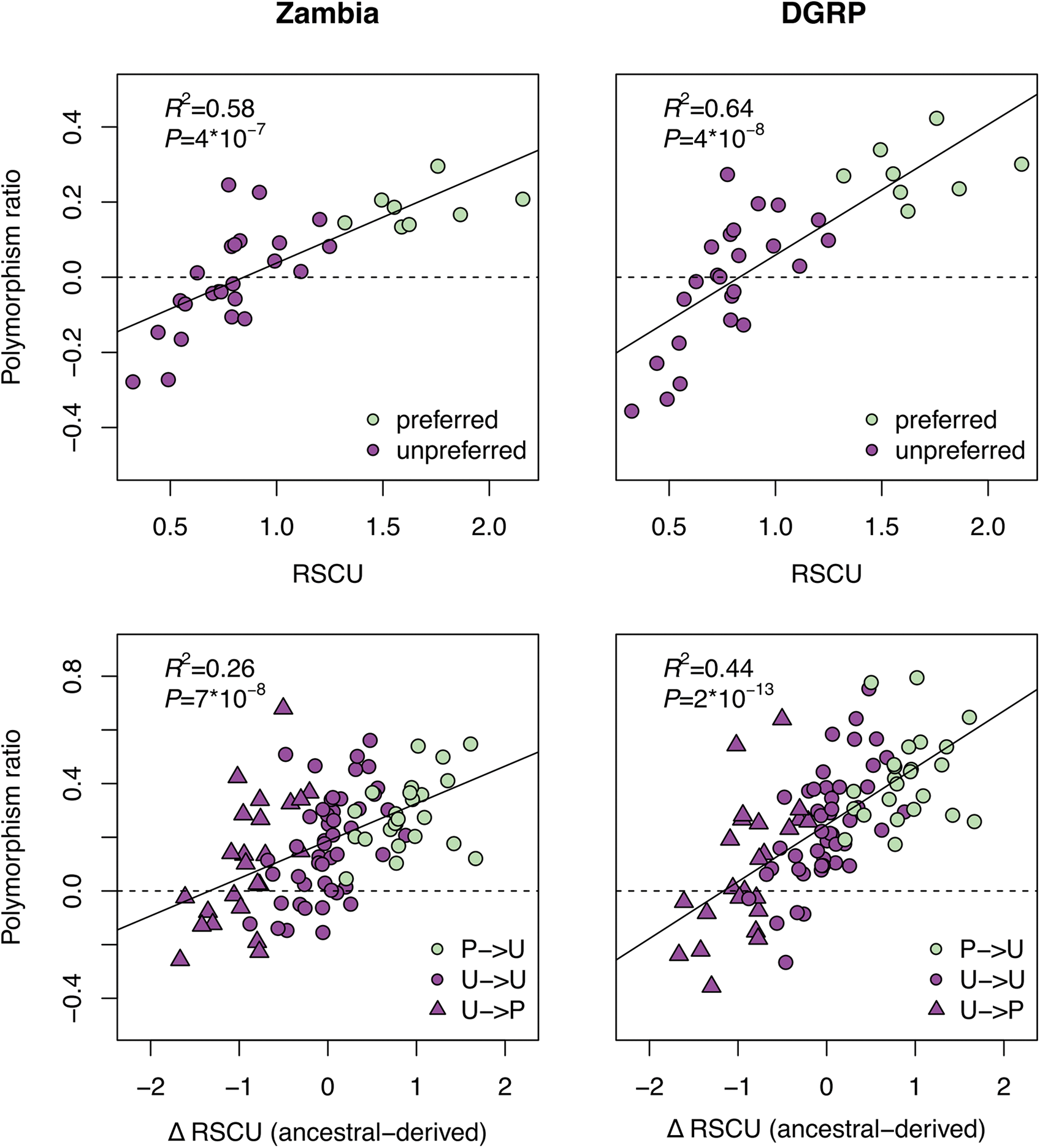
Top: Polymorphism ratio for each codon as a function of the level of bias for the codon (relative synonymous codon usage: RSCU) (median over 200 matched controls). Bottom: Polymorphism ratio for each ancestral/derived codon pair as a function of the change in RSCU.

### 3.6 More selection on synonymous sites due to CUB than due to any other process

Several processes other than those related to CUB have also been hypothesized to act on synonymous sites. In order to assess the relative importance of various processes driving the observed selection on synonymous sites, we tested several putatively functional classes of sites for enrichment of purifying selection. In addition to preferred codons, we tested transcription factor (TF) bound regions, alternatively spliced genes, RNA binding protein (RBP) bound regions, splice junctions and high ribosomal occupancy regions. We calculated the polymorphism ratio for each functional class and the corresponding dataset excluding the functional class (exclusion dataset). We found a significantly greater polymorphism ratio not only for preferred codons but also for alternatively spliced genes, spliceosome bound regions and TF bound regions in both the Zambia and the DGRP populations (Figure 5, Supplementary Figure 6).

**Figure 5:**
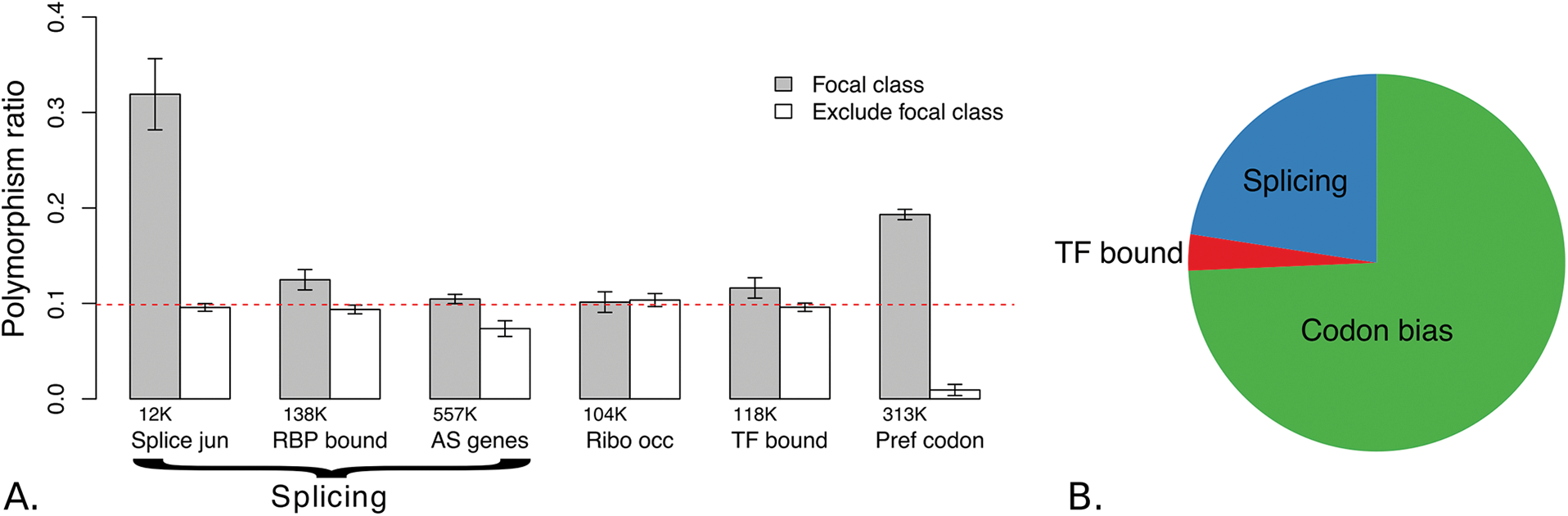
A) Proportion of sites under strong selection as measured by the polymorphism ratio for each class of site (grey) and the dataset excluding the focal class sites (white). The number of sites in a focal class is listed below the corresponding bar. The red dashed line is the polymorphism ratio for the full dataset (Zambia). Error bars represent two standard error. B) Relative proportion of synonymous sites under strong purifying selection due to slicing, codon bias, or being transcription factor (TF) bound.

In order to compare the relative contributions of each functional class to strong purifying selection, we estimated the number of sites expected to be under strong purifying selection as a result of a particular functional class (see Methods). We combined the three splicing-related classes (alternatively spliced genes, spliceosome bound regions, and splice junctions) into one “splicing” dataset and compared this to the set of sites not covered by any of these categories. This left us with three general groups: codon bias, splicing, and transcription factor binding. In the Zambia dataset we found 150K sites under strong purifying selection associated with codon bias, 38K with splicing, and 4K with transcription factor binding (Figure 5). The DGRP dataset showed similar trends: 217K sites under strong purifying selection associated with codon bias, 100K with splicing, and 13K with transcription factor binding. In summary, we found that codon bias explained the greatest number of 4D sites under purifying selection, representing approximately twice as many sites as splicing.

We also measured the polymorphism ratio for the sites least likely to be under selection. We excluded the two largest contributors to selection on synonymous sites, preferred codons and alternatively spliced genes. The set of unpreferred codons in non-alternatively spliced genes consisted of 137K sites in Zambia and 158K sites in DGRP, and represented the 4D sites least likely to be under strong selection. Interestingly, we found that this set of 4D sites had more polymorphism than their SI matched control set (negative polymorphism ratio), indicating greater purifying selection in short introns and/or the presence of positive selection on these 4D sites.

### 3.7 Features of strong selection on CUB

Preferred codons may be under greater purifying selection in some genes than in others. We asked if a greater proportion of preferred codons were under strong purifying selection in genes with high codon bias compared to genes with low codon bias. One measure of the amount codon bias in a gene is the frequency of preferred codons (FOP). We calculated FOP per gene and asked if, as expected, there was a stronger signal of purifying selection on 4D sites in genes with higher FOP. We found a trend towards a larger polymorphism ratio for 4D sites in high FOP genes (Zambia: 0.103; DGRP: 0.150) compared with low FOP genes (Zambia: 0.087; DGRP: 0.131) albeit the trend is not significant (Zambia: *t*-test *P* = 0.19; DGRP: *t*-test *P* = 0.26; Supplementary Figure S4).

We then evaluated the patterns of CUB-associated polymorphism by grouping 4D sites into three categories: preferred, unpreferred with mutations to another unpreferred state, unpreferred with mutations to the preferred state. We found no trend of polymorphism ratio verses FOP for preferred codons, indicating that a similar proportion of preferred codons were under strong selection in genes with low overall biased codon usage compared with genes with high bias and consequently, that a larger *number* of preferred codons in high FOP genes are subject to strong selection (Figure 6). Interestingly, we found a pattern of negative polymorphism ratios for unpreferred codons specifically in high FOP genes, which was particularly pronounced for sites that were ancestrally unpreferred with derived preferred mutations. This pattern was much stronger in high FOP genes than low FOP genes (Zambia: *t*-test *P* = 0.02; DGRP: *t*-test *P* = 3 ∗ 10^−5^). Note that these negative polymorphism ratios at unpreferred codons lead to lower polymorphism ratios in high FOP genes than would be expected given the larger number of preferred codons subject to strong selection in such genes (Supplementary Figure S4). These patterns overall are consistent with stronger selection in favor of preferred codons in high FOP genes.

**Figure 6:**
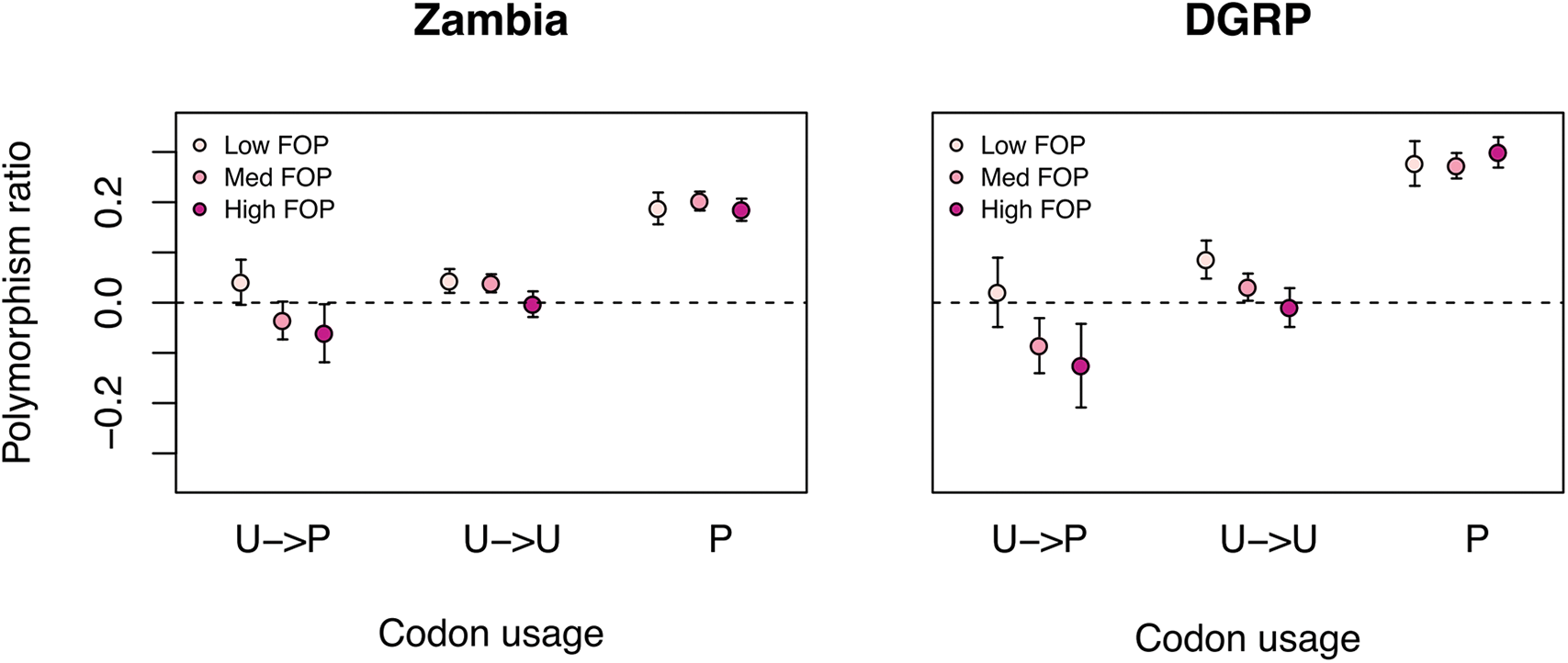
Polymorphism ratio by class of codon preference and the frequency of preferred codons (FOP). FOP groups are classified into low (lowest quartile), medium (middle two quartiles), and high (top quartile). U: unpreferred; P: preferred. Error bars are 2 standard error.

Codon bias has also been shown to vary depending on the location in the gene. We first asked if preferred codons vary in the amount of strong purifying selection that they are under as a function of the location in the exon. We measured the polymorphism ratio for each class codon preference at the start (1st quartile) of an exon, the middle of an exon (2nd and 3rd quartile) or the end of an exon (4th quartile). In preferred codons we found a trend toward increased polymorphism ratio at the start and the end of exons, compared with the middle of the exons (Figure 7; *t*-test start > middle: Zambia *P* = 0.1, DGRP *P* = 0.01; *t*-test end > middle: Zambia *P* = 8 ∗ 10^−5^, DGRP *P* = 0.03). However, this pattern was also observed in unpreferred codons (*t*-test start > middle: Zambia *P* = 0.05, DGRP *P* = 0.04; t-test end > middle: Zambia *P* = 0.06, DGRP *P* = 0.5), indicating that this effect may be unrelated to CUB. Alternatively, this could be a result of purifying selection on synonymous sites important for splicing. We next assessed polymorphism ratio as a function of the exon position along the gene (either first exon, last exon, intermediate exons, or exons of single-exon genes). No consistent patterns were observed with location of the exon (Figure 7).

**Figure 7:**
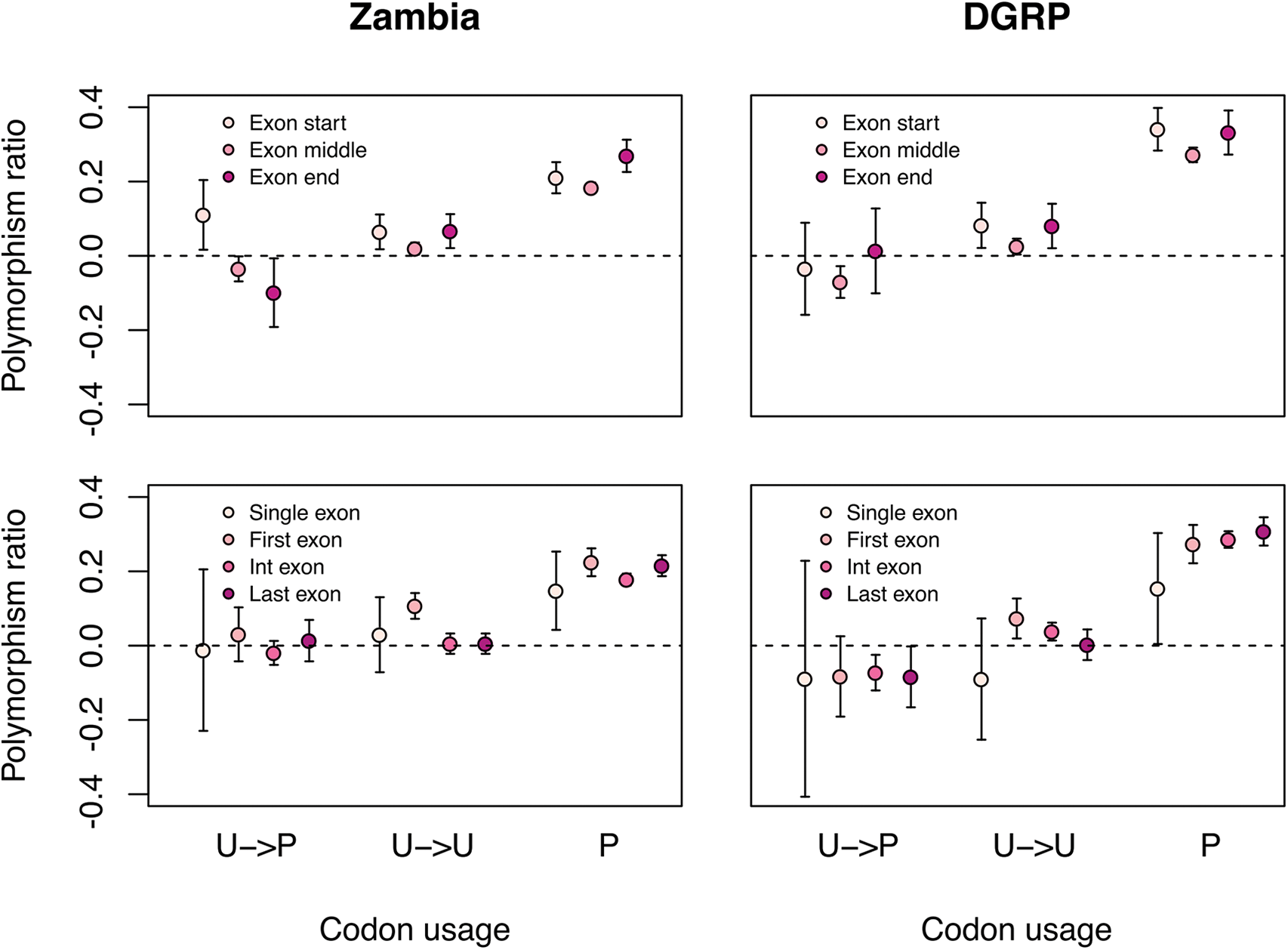
Polymorphism ratio by class of codon preference and the position along the exon (top) or the exon position along the gene (bottom). U: unpreferred; P: preferred. Error bars are 2 standard error.

## 4 Discussion

### 4.1 Strong and weak purifying selection on CUB

We find evidence that selection on CUB is not limited to weak selection, and find that ∼ 20% of 4D sites in preferred codons are under strong purifying selection. Our study builds on methodology developed in Lawrie *et al.* 2013, recapitulating their major result of strong purifying selection on synonymous sites and extending the analysis to identify functional associations. We were able to gain a finer view by 1) use of multiple, deeply sampled datasets, 2) ancestral polarization of alleles, and 3) strict filtering of sites with low quality or near indels to reduce noise. Although Lawrie *et al.* 2013 found evidence of strong selection on 4D sites, the underlying processes examined could not account for this signal. While our finding of strong purifying selection on 4D sites is consistent with Lawrie *et al.* 2013, our study finds that CUB accounts for the majority of this selection, and, in conjunction with splicing, can fully explain the patterns of polymorphism. In addition to finding that many 4D sites are subject to strong selection, we also find evidence that a substantial proportion of 4D sites are under weak purifying selection on CUB, which is consistent with the signal of weak selection previously observed in *D. melanogaster* (Zeng & Charlesworth 2009, 2010, Campos *et al.* 2013).

For methodological reasons, many previous methods identified only weak selection on CUB. Strong purifying selection is not detectable with methods that use only the polymorphic SFS without sufficiently high depth of population sequencing (Zeng & Charlesworth 2009, 2010, Campos *et al.* 2013) or methods that incorporate polymorphism-level, but assume a distribution of fitness effects (DFE) that is biased towards weak-selection (eg. gamma distribution: Andolfatto *et al.* 2011). In our analysis we use the polymorphism-level and SFS to make point estimates of selection strengths. We detect both a peak of selection coefficients at *N_e_s* = −1 as well as at *N_e_s* = −22 in the set of preferred codons (DGRP: *N_e_s* = −1, *N_e_s* = −66). In reality, selection coefficients have a distribution, which we have represented with either one or two selection masses. The use of point estimates to represent the DFE is robust to a range of real underlying DFEs (Kousathanas & Keightley 2013), allowing us to detect selection occurring at both the weak and the strong range of selection coefficients.

### 4.2 Polymorphism ratio correlates with the level of CUB per codon and per gene

Since we control for mutation rate and local determinants of polymorphism, such as linked selection and recombination, we can use polymorphism-level information alone to measure strong purifying selection. We find that the polymorphism ratio of the SI to 4D sites is a good proxy for the proportion of sites under strong purifying selection, as evidenced by the relationship between polymorphism ratio and both the ML estimates of selection and the level of phylogenetic conservation. We find that the estimated proportion of sites under strong selection is strongly associated with the extent of CUB, as measured by the relative synonymous codon usage (RSCU). In addition, the change in RSCU from ancestral to derived correlates with the proportion of sites under strong selection. These results further support our conclusion of strong purifying selection on CUB.

It is well established that certain genes, particularly those with high expression, tend to have a greater proportion of preferred codons (Gouy 1982; Bulmer 1991; Novoa & Ribas de Pouplana 2012). We measured polymorphism ratio for sites in genes of low, medium, and high frequencies of preferred codons (FOP). From this analysis we have three major findings: 1) the proportion of preferred codons under strong purifying selection is relatively constant across genes (Figure 6), 2) there is evidence for increased positive selection for derived preferred mutations in high FOP genes (Figure 6), and 3) the contribution of excess 4D polymorphism, putatively associated with positive selection, in high FOP genes is a example of how the polymorphism ratio measure of strong purifying selection can be dampened by positive selection. To more fully articulate the third point, polymorphism ratio in high FOP genes is the combination of two competing processes, the higher proportion of preferred codons increasing the polymorphism ratio and the stronger positive selection for derived preferred codons reducing the polymorphism ratio. The fact that the increase in polymorphism ratio between low and high FOP genes is less than expected (Supplementary Figure S4) is a demonstration of cryptic CUB-associated purifying selection in the high FOP genes (due to the increased level of positive selection on CUB). We also notice negative polymorphism ratios in the RSCU analysis, where we find strongly negative polymorphism ratios for codons that we would expect to be under the greatest amount of positive selection, i.e., codons with highly unpreferred ancestral states and highly preferred derived mutations.

### 4.3 Selection on other functional classes

We find splicing to be the second-most important process underlying purifying selection on synonymous sites. We tested three classes of sites putatively enriched for selection due to splicing: alternatively spliced genes, spliceosome-bound regions and splice junctions. Although alternatively spliced genes explain the greatest amount of selection on synonymous sites (∼90K sites), owing to the large number of sites in alternatively spliced genes, we find that splice junctions have the greatest proportion of sites under selection (∼ 45% under strong purifying selection), followed by spliceosome-bound regions. Splicing is known to be a critical function for proper development and function of an organism.

There is also evidence for an enrichment of strong selection in transcription factor-bound 4D sites. We estimate that ∼3K 4D sites are under strong selection due to transcription factor binding. To identify transcription factor bound sites we used ChIP-seq experiments targeted at 16 different transcription factors. With a larger breadth of transcription factor binding data, 4D sites in transcription factor-bound regions may prove to be under a greater amount of selection than we can detect here.

We find that codon bias, splicing, and transcription factor binding are sufficient for explaining the polymorphism differences between 4D and SI control sites, indicating that these processes also explain the bulk of strong purifying selection acting on synonymous sites. However, it is important to note that our measures are only correlative with the functional class being tested, such that we cannot say that these processes directly underlie the selection. In addition, there are likely multiple other processes acting on synonymous variants that we have not included. Other processes that have been shown or hypothesized to act on 4D sites include transcriptional regulation (Newman *et al.* 2016) and RNA transcript stability (Presnyak *et al.* 2015). Given the explanatory power of our results, we suggest that these other processes are either less affected by synonymous variation or that they are correlated with the processes already tested.

### 4.4 Controlling for linked selection and mutation rate

One caveat to our polymorphism-level based method of estimating selection is that multiple processes can reduce the observed level of polymorphism of a site. These include linked selection, low recombination rate, a reduced mutation rate or selection on the site itself. In order to isolate the effects of selection on 4D sites, we ensured that each 4D site was experiencing the same local environment of linked selection and recombination rate and the same mutation rate as its matched SI control. We found that with an increasing distance of up to 1000bp from the focal 4D site to its SI control there was no systematic change in polymorphism in the SI control, indicating that the matched controls were under a sufficiently similar amount of linked selection (Supplementary Figure 1). This local matching also ensures equivalent recombination rates, which can affect polymorphism. In order to account for mutational differences, we required that matched controls had the same 3bp mutational context as the 4D sites. In *Drosophila*, there is a significant effect of 3bp context on mutation rate (Sharp & Agrawal 2016). For polymorphic sites we used the ancestral allele for matching (polarized from *D. simulans*, where possible), providing a more appropriate match than if we had not polarized by ancestral state. Since we match locally (< 1000bp), 4D sites and their matched controls will also be subject to the same local mutation rate effects, such as GC content. In addition to taking measures to control for mutation rate, we observe that our estimates of purifying selection correlate with the putative functionality of a class of sites, such as preferred codons, splice junctions, RBP bound regions, and alternatively spliced genes, supporting the claim that our results reflect the action of selection.

### 4.5 Our selection estimates may be conservative

Our estimates of purifying selection on 4D sites may be conservative, underestimating the true amount of selection on 4D sites. This could be the case if there was any constraint on the SI controls or if there was positive selection on the 4D sites themselves. There were two methodological decisions that may have contributed to constraint in short introns. Both (Halligan & Keightley 2006) and (Parsch *et al.* 2010) found that short introns (< 65bp and < 120bp, respectively) have the least constraint on bases 8-30. As we included a larger portion of the intron, it is possible that we have also included SI sites under a greater amount of conservation. We also excluded regions surrounding indels (10bp on either side) in order to reduce false polymorphisms due to mis-mapping. This more strongly affects short introns (as they are more permissive to indels than coding regions) and will select for more conserved SI regions. We also find evidence supporting positive selection on 4D sites, where 4D sites in ancestrally unpreferred codons with a derived preferred allele actually have an excess of polymorphism compared to the SI controls.

### 4.6 New model of CUB

Our finding that selection on CUB ranges from weak to strong directly contradicts the standard Li-Bulmer model of selection on CUB. The Li-Bulmer model assumes a constant selection coefficient for a codon and, given the intermediate proportion of preferred codons observed in many species, predicts that selection on CUB is weak (Bulmer 1991, Li 1987). This prediction may have contributed to the prevalence of methods that are biased towards the detection of weak selection. However, the Li-Bulmer model has not always agreed with the data. First, since population sizes vary by several orders of magnitude across species, the selection coefficient would have to vary inversely by several orders of magnitude as well in order to result in the observed intermediate levels of CUB (Hershberg & Petrov 2008). There is no intuitive reason to think that the selection coefficient would be inversely related to the population size, or that it should vary by several orders of magnitude. Second, if selection is weak, there should be more CUB in high recombination rate regions. This prediction comes from the increased effect of Hill-Robertson interference (linked selection) in low recombination rate regions (Felsenstein 1974). While there is some evidence for a correlation between CUB and recombination rate in *D. melanogaster* (Kliman & Hey 1993; Campos *et al.* 2012), this is not true for the *D. melanogaster* X chromosome (Singh 2005; Campos *et al.* 2013), and the correlations that have been found can be explained by mutation rate (Marais 2001). Third, there is experimental evidence that changes in one or more synonymous codons can have large phenotypic effects, suggesting that selection on CUB is not always weak (Zhou *et al.* 1999, Carlini & Stephan 2003, 2004).

We propose a new model where the strength of selection per codon varies from non-existent to strong within a gene, with the level of CUB in a gene set primarily by the distribution of selection coefficients across sites. Genes that have high CUB under our model would have more sites subject to strong selection in favor of preferred codons compared to genes with low CUB, as we in fact see in the data. This eliminates the problem of setting the proportion of preferred codons by fine-tuning the strength of selection at all preferred sites to a particular value of *s* ∼ 1/*N_e_* under the Li-Bulmer model. In addition, under our model, a substantial proportion of preferred codons is subject to such strong purifying selection (*s* >> 1/*N*_*e*_), that reduction in effective population size by orders of magnitude due either to demographic shifts or modulation in the strength of genetic draft would still not abolish CUB, as many preferred sites would still remain subject to strong selection (*s* > 1/*N*_*e*_). At the other extreme, a substantial increase in effective population size would not generate complete CUB as many preferred sites may not be subject to purifying selection at all.

If this model is correct, the key question that remains is what determines whether a particular synonymous site is subject to strong, weak, or no selection in favor of preferred codons. Specifically, the sites under very strong selection might play a disproportionately important role by, for example, being essential for cotranslational folding, transcription, RNA stability, translational efficiency or translational accuracy. This would suggest that the location of such synonymous sites should be largely conserved across species, as we in fact detect to some extent by showing a correlation between polymorphism ratio and phylogenetic constraint in the *Drosophila* genus (Figure 2).

### 4.7 Conclusions

We find evidence that codon usage bias is under a substantial amount of purifying selection in *D. melanogaster*, and that this is not limited to weak selection. Our finding that there is a distribution of fitness effects for CUB, ranging from weak to strong selection, argues against the Li-Bulmer model predicting constant weak selection. By dismissing this model, we resolve the contradiction between the intermediate frequencies of preferred codons observed in most species and the population-size independence of said frequencies. We also reconcile the observations that changes in synonymous codons can have large phenotypic effects, but that genomic methods have identified only weak selection. We suggest that the reasons previous studies did not find evidence for strong selection on CUB are methodological. Our use of a test that includes the polymorphism-level, while controlling for mutation rate and linked selection, provides sufficient power for identifying strong purifying selection. While this study was performed in *Drosophila*, the importance of a new model of CUB is general, as both codon bias and the assumption of constant weak selection is widespread. Further, this study underscores the importance of CUB, and of synonymous variation in general, to the fitness of an organism, and opens research directions to further understand this phenomenon.

## 5 Methods

### 5.1 Sequence data

We used sequence data from two *D. melanogaster* populations, one from North America (DGRP Freeze 2), consisting of 200 inbred lines (Mackay *et al.* 2012), and one from Africa (Zambia), consisting of 197 haploid embryos (Lack *et al.* 2015), downloaded from the Drosophila Genome Nexus (http://www.johnpool.net/genomes.html). To reduce the effect of sequencing and mapping error, for each individual we filtered out all sites with low mapping quality (MAPQ < 20) and that were within 10bp of an indel. Per population we down-sampled sites to a uniform coverage of 160X and excluded sites with less than 160X coverage. We considered only the four major autosomal chromosome arms because of systematic differences between *D. melanogaster* autosomes and X chromosomes (Singh *et al.* 2005) and polarized polymorphic sites by identifying the ancestral state as the allele found in the *D. simulans* v2 reference genome (Hu *et al.* 2013). We used the *D. melanogaster* reference allele for cases where the ancestry was ambiguous, either because there was no direct *D. simulans* alignment or because neither allele was present in *D. simulans*. Fourfold degenerate synonymous (4D) sites and intronic regions were identified from Flybase annotations (release 5.5; www.flybase.org). The total number of 4D sites in our two datasets was 1,976,830 for DGRP and 1,862,290 for Zambia. We classified short introns (SI) as introns less than 86bp in length and excluded the first and last 8bp of each intron, as these regions are known to be under constraint (Haddrill *et al.* 2005; Halligan & Keightley 2006, Clemente & Vogl 2012). The total number of SI sites was 550,587 for DGRP and 446,462 for Zambia.

We created the SI control dataset by matching each 4D site to a SI site. To control for mutation rate differences between 4D sites and their matched controls, we required each matched SI site to have the same ancestral allele and the same neighboring nucleotides (3bp context) as the 4D site. We matched blind to the direction or strand (i.e., matching with the forward, reverse, reverse complement, or complement SI sequence). To control for the effect of linked selection on the level of 4D polymorphism, we also required each matched SI site to be within 1000bp of the 4D site, such that SI control would be subject to the same linked selective pressure from nonsynonymous sites as the 4D sites. We found 1000bp to be a sufficiently small distance, as we found no significant correlation between SI polymorphism and distance between the 4D sites and the matched intron over the range of 0 to 1000bp (Supplementary Figure 1). We produced 200 matched sets, each with the same 871,218 DGRP or 754,503 Zambia 4D sites, and an average of 288K SI sites for DGRP and 244K SI sites for Zambia (a given SI site may be matched to multiple 4D sites).

### 5.2 Maximum-likelihood estimation of selection parameters from SFS

We employed a variation of the site frequency spectra (SFS) method described in Lawrie *et al.* 2013. The method uses both SNP density and frequency information of SFS to calculate the distribution of fitness effects (DFE) for a test set of sites given a “neutral” reference - in this case, the DFE for 4D synonymous sites with SI sites as the reference. While during bootstrapping SNPs are polarized for ancestral state, for the purposes of maximum-likelihood estimation, the spectra are folded - which restricts the analysis to purifying selection. The DFE itself is modeled as a categorical distribution where the program estimates selection coefficients (*N_e_s*) and the percentages of sites (*f*) evolving under those selection coefficients for a predetermined number and type of selection categories. This has the advantage of not assuming a particular distribution shape such as gamma or lognormal, but comes at the cost of additional free parameters per additional categories. For example, a three category model which has a neutral class (*f*_0_) + a weak selection class (*f*_*W*_, 0 > *N*_*e*_*s*_*W*_ > −10) + a strong selection class (*f*_*S*_, −10 > *N*_*e*_*s*_*S*_ > −*inf*) requires 4 free parameters to fully describe it (*f*_0_ = 1 − *f*_*W*_ − *f*_*S*_, *N_e_s*_0_ = 0). The method also estimates the scaled mutation rate, *θ* (*N_e_µ*), for the SI spectra.

Demography, linked selection, and other forces affecting both 4D and SI sites, can skew the spectra and bias the estimation of the above DFE parameters. To compensate, we used frequency-dependent correction factors, *α*_*x*_, which adjusts the probability of seeing a site with a SNP at frequency of *x* in the sample - *p*(*x*|model) (see Lawrie *et al.* 2013). The likelihood (*λ*) of the SFS under the model’s framework is shown below:

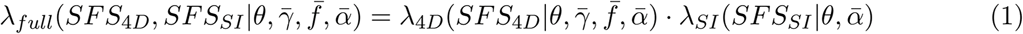

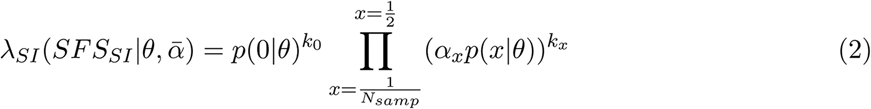

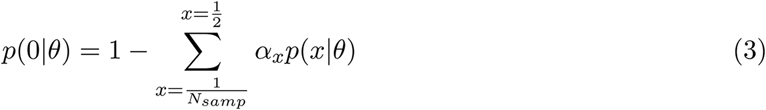

where *α*_1_/*N*_*samp*_ = 1, *N*_*samp*_ = number of frequencies in the population sample, and where *α*_0_ = 1 and *k*_*x*_ is the number of polymorphic sites at frequency *x* in the *SFS*_*SI*_. Matlab code for ML testing is available on Github.

#### 5.2.1 Model adjustment for demography

Deviations of the putatively neutral SI SFS from the theoretical neutral SFS are expected to exist due to an organism’s demographic history. To account for this deviation of the SI SFS from the theoretical neutral, we preformed a maximum likelihood fit of offsets (alpha values) for each allele frequency bin. Allele frequency bins were divided, according to a power law, into 6 separate bins. (Supplementary Table 1). We found a good fit of the demography-corrected SFS to the SI SFS (i.e. two distributions are not significantly different, KS test *P* = 1).

#### 5.2.2 Model parameters

We tested five different ML models: 1) neutral, 2) neutral + lethal, 3) neutral + 1 selection coefficient, 4) neutral + selection + lethal, and 5) neutral + 2 selection coefficients. We ran the ML estimation both with and without (SFS only) polymorphism-level data. The neutral + 2 selection coefficients model requires a parameter that is the boundary condition between weak and strong selection classes. We tested a broad range of boundary conditions and found *Ns* = −10 to permit all maximum likelihood peaks to be reached. The ML test required seed values for selection strength, selection proportion, lethal proportion and theta. After a rigorous search of the parameter space, we identified the highest likelihood model. To calculate 95% confidence intervals, we performed a rank bootstrap, sampling with replacement each of the 200 matched 4D and SI datasets, performing our maximum-likelihood estimate of selection and using the 5th and the 195th rank values for each maximum-likelihood score, proportion of selection, and strength of selection. To determine the best fit model, we performed a chi-squared likelihood ratio test of the maximum-likelihood scores.

#### 5.2.3 Power analysis

In order to assess our power in differentiating strong selection from a lethal class or 4D/SI mutational differences, we performed power analyses of our level + shape maximum likelihood method of selection estimation. We did this by creating a theoretical SFS’s for a range of selection strengths and proportions, and for theta values reflecting those of the DGRP (0.01) and Zambia (0.035) populations and estimating selection for this using a theoretical neutral reference with the same theta value and number of sites. We performed a chi-squared likelihood ratio test with one degree of freedom comparing the 2-category selection model (neutral + one selection class) with the neutral + lethal model. This differs from our main analysis in that we did not perform bootstrap replicates and did not calculate the corresponding rank bootstrap confidence interval. This analysis demonstrates how an increasing number of SNPs, increasing polymorphism level (eg. larger theta), and a greater proportion of sites under selection increase our power to distinguish strong selection from lethality/mutational differences.

### 5.3 Polymorphism ratio estimate of strong selection

In order to make a precise estimate of purifying selection using our sfs-based maximum likelihood method, we require a large number of sites (> 100K). When we have few sites, we can use alternative methods for estimating purifying selection. One proxy for the amount of strong purifying selection is the depletion of polymorphism in a selected class compared with a neutral class. We quantified this depletion as the “polymorphism ratio”:

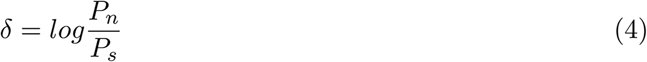

where *P* is polymorphism, *s* is the selected class (4D sites) and *n* is the neutral class (SI sites). This statistic is positive when polymorphism is greater in SI sites and negative when polymorphism is greater in 4D sites. For all analyses we used the median polymorphism ratio of 200 matched control sets. We found a strong correlation between the polymorphism ratio and the estimated proportion of sites under strong selection (*R*^2^ = 0.95; Figure S3).

We estimated the number of sites expected to be under strong purifying selection as a result of a particular functional class (*N*_*sel*_) as:

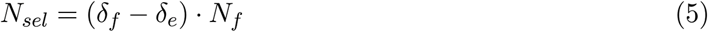

where *f* is the focal dataset, *e* is dataset excluding the focal sites and *N*_*f*_ is the number of sites in the focal dataset.

### 5.4 Identification of putatively functional regions

#### 5.4.1 Codon Bias

We calculated the relative synonymous codon usage (RSCU) for each codon as the observed frequency of a codon in the dataset divided by the expected usage if all four codons were used equally (0.25) (Sharp & Li 1986). We classified each 4D site as being in a preferred (highest RSCU for the amino acid) or unpreferred codon (lowest three RSCU’s for the amino acid). The amino acids and their respective preferred codons are as follows: alanine GCC, glycine GGC, leucine CTG, proline CCC, threonine ACC, and valine GTG. For polymorphic 4D sites we used the ancestral allele to designate the codon. We identified at total of 850,973 (509,997 with SI controls) and 794,471 (458,356 with SI controls) 4D sites in preferred codons for DGRP and Zambia, respectively.

For each codon-changing 4D mutation, we measured the change in RSCU from the ancestral to the derived codon. We then examined the relationship between RSCU change and polymorphism ratio. In order to appropriately calculate the polymorphism ratio for each codon change, we matched 4D sites to SI sites with the same possible states. For example, for the class of 4D sites of an ancestral “CCC” proline codon and a derived “CCA” proline codon, we matched the 4D proline “C” monomorphic sites and derived “A” polymorphic sites to SI “C” monomorphic sites and derived “A” polymorphic sites (or the complement), as well as matching for distance and mutational context.

#### 5.4.2 Transcription Factor Binding Sites

We used modEncode chromatin immunoprecipitation sequencing (ChIP-seq) experiments to assess the contribution of transcription factor binding sites to the signal of purifying selection on synonymous sites. This dataset represents 25 experiments, testing 15 transcription factor targets (antibodies: odg-GFP, anti-trem, Sin3A-RC, Su(var)3-9, KW4-PCL-D2, KW3-D-D2, KW3-Trl-D2, bon (GP37), HP1 antibody (ab24726), HP1-Covance, KW4-Hr39-D1, KW3-Kr-D2, KW3-CG8478- D1, KW3-hkb-D1, KNI-D2,KW3-Trl-D2; modENCODE submissions 3229, 3230, 3232, 3234, 3237, 3238, 3239, 3240, 3241, 3242, 3243, 3245, 3390, 3391, 3392, 3393, 3394, 3395, 3396, 3398, 3399, 3400, 3401, 3402, 3403). We consider a “transcription factor bound region” any region with evidence for TFB in any of the non-control experiments (minimum binding score: 50). We identified a total of 294,703 (173,334 with SI controls) and 289726 (164,842 with SI controls) transcription factor bound 4D sites for DGRP and Zambia, respectively.

#### 5.4.3 Spliceosome binding

We used modEncode RNA immunoprecipitation sequencing (RIP-seq) experiments targeting putative spliceosome proteins to assess the contribution of spliceosome binding to the signal of purifying selection (http://intermine.modencode.org). The experiments tested for RNA-protein binding of a total of 30 putative splicing proteins. We considered a region to be bound if it had a binding score of 5 of greater in any of the experiments. This left a total of 321,290 (204,901 with SI controls) and 316,740 (194,046 with SI controls) spliceosome-bound 4D sites for DGRP and Zambia, respectively.

#### 5.4.4 Alternative splicing

We distinguished between genes with and genes without alternative splicing using the analysis in Brown *et al.* 2014. We considered any gene with more than one transcript as alternatively spliced. We found a total of 1,196,063 (864,846 with SI controls) and 1,136,535 (792,445 with SI controls) 4D sites in alternatively spliced genes for DGRP and Zambia, respectively.

#### 5.4.5 Splice junctions

We used the splice junctions identified by Brooks *et al.* 2015. We found 18410 and 17528 4D sites in splice junctions for DGRP and Zambia, respectively.

#### 5.4.6 Ribosomal occupancy

We estimated ribosomal occupancy using the ribosomal profiling experiments conducted by Dunn *et al.* 2013. We first normalized each pooled experiment files (GEO accession GSE49197) by dividing the number of counts in each region by the total number of counts across regions. All regions with zero counts for either the footprinting or expression experiments were excluded. We estimated translational efficiency by dividing the normalized ribosomal footprint values by the normalized expression values (for each DNA strand separately). The top and bottom 1 percentile of ribosomal occupancy scores were omitted from downstream analysis, leaving translational efficiency scores for 1,391,585 4D sites. We divided these sites into three categories, high, medium, and low ribosomal occupancy, based on the lowest third, the middle third, and the top third of values, respectively.

#### 5.4.7 Frequency of preferred codons

We calculated the frequency of preferred codons (FOP) per gene. As before, preferred codons were defined as the most frequent codon for a given amino acid. The FOP was calculated with our 4D datasets, such that codons that did not appear in our datasets (eg. those without 4D sites) did not contribute to the FOP calculation. Sites were classified as being in genes with either low (bottom quartile), medium (middle two quantile), or high (top quantile) FOP. The average proportion of preferred codons for sites in low, medium, and high FOP genes was 28, 42, and 54 percent, respectively.

### 5.5 Conservation scores

We calculated the level of conservation of each 4D site across a 10-species *Drosophila* phylogeny that excluded the focal species, *D. melanogaster*. The PRANK multiple sequence alignments of the 10 species (*D. simulans, D. sechellia, D. yakuba, D. erecta, D. ananassae, D. pseudoobscura, D. persimilis, D. virilis, D. mojavensis, D. grimshawi*) were generously provided by Dr. Sandeep Venkataram. We calculated the probability of conservation for each 4D site using the *phyloP* function of the PHAST software (method=LRT) (Cooper *et al.* 2005). Given the size of the phylogeny, the highest significance score for conservation was *P* = 0.15. Thus, we identified a conserved site as one with a phyloP *P* < 0.2.

## 6 Supplementary Figures

**Figure S1:**
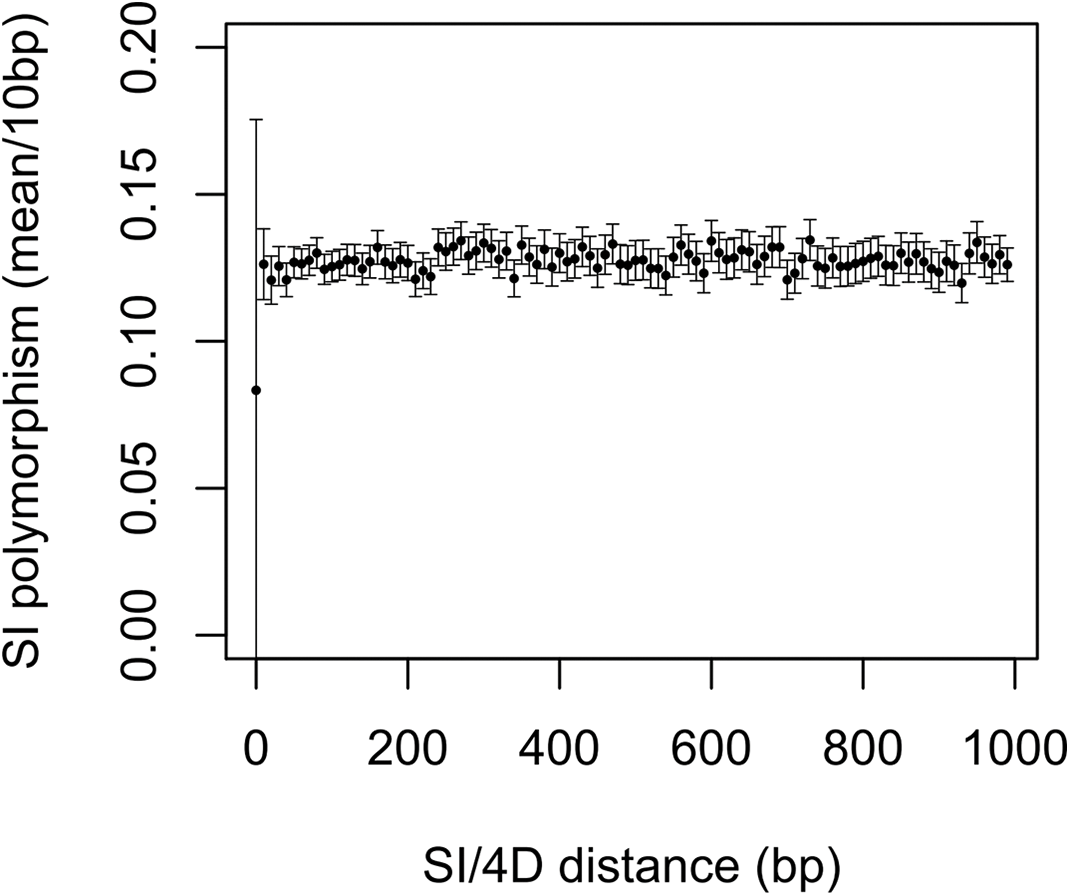
Short intron polymorphism as a function of distance from the 4D site.

**Figure S2:**
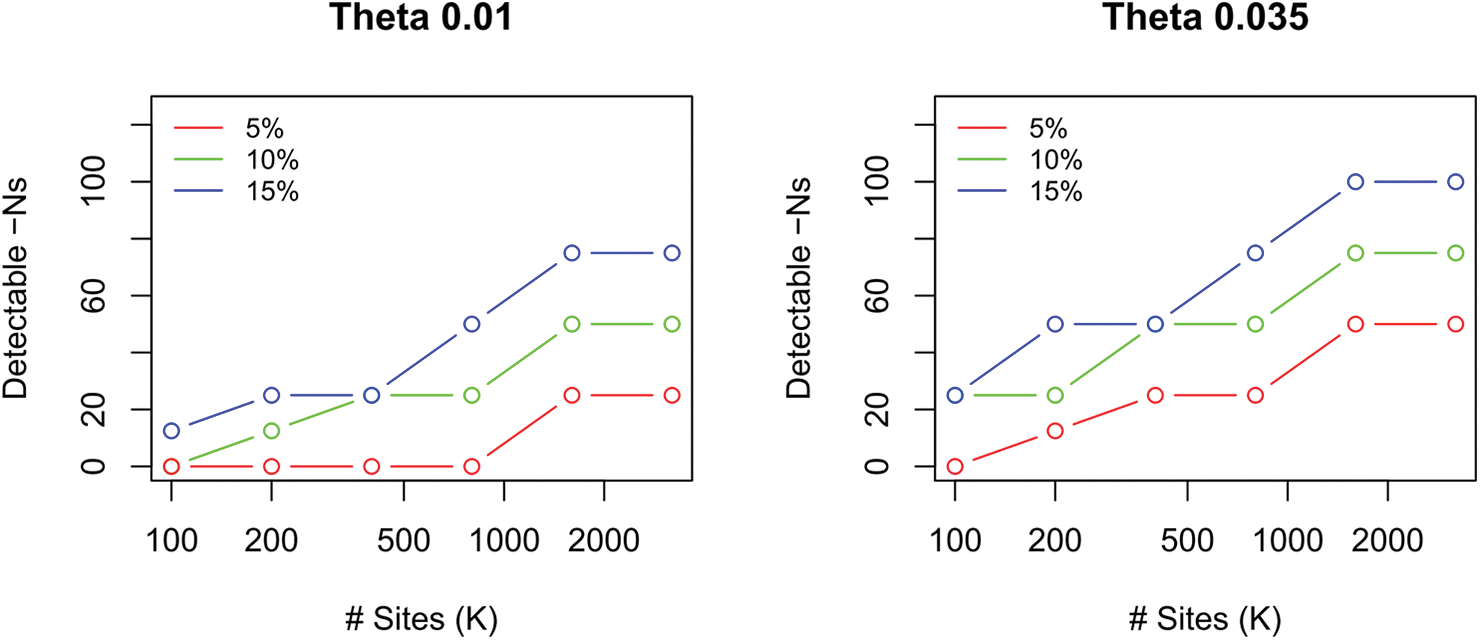
The maximum strength of selection detectable (significantly distinguishable from lethality) as a function of the number of sites analyzed (in 1000’s of sites), for a range of proportions of sites under selection (red: 5%, green: 10%, blue: 15%).

**Figure S3:**
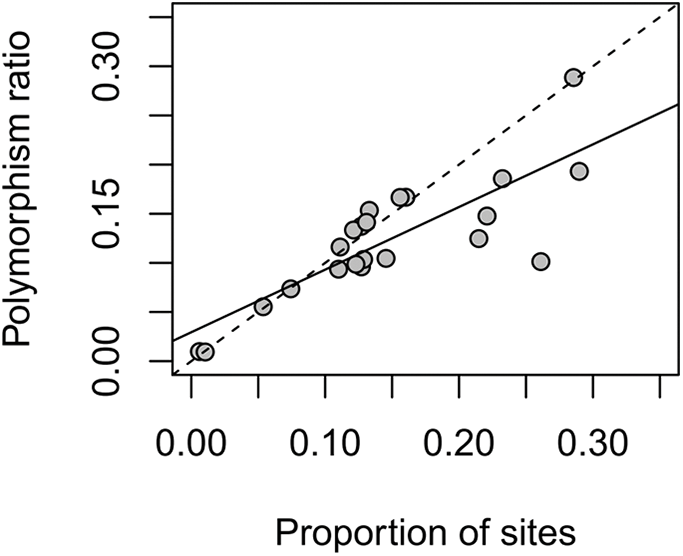
Correlation between the polymorphism ratio and the proportion of sites estimated to be under selection (1 selected class model). Each point represents a different subset of the Zamiba or DGRP datasets (functional classes tested). Solid line is a linear regression (*R*^2^ = 0.65).

**Figure S4:**
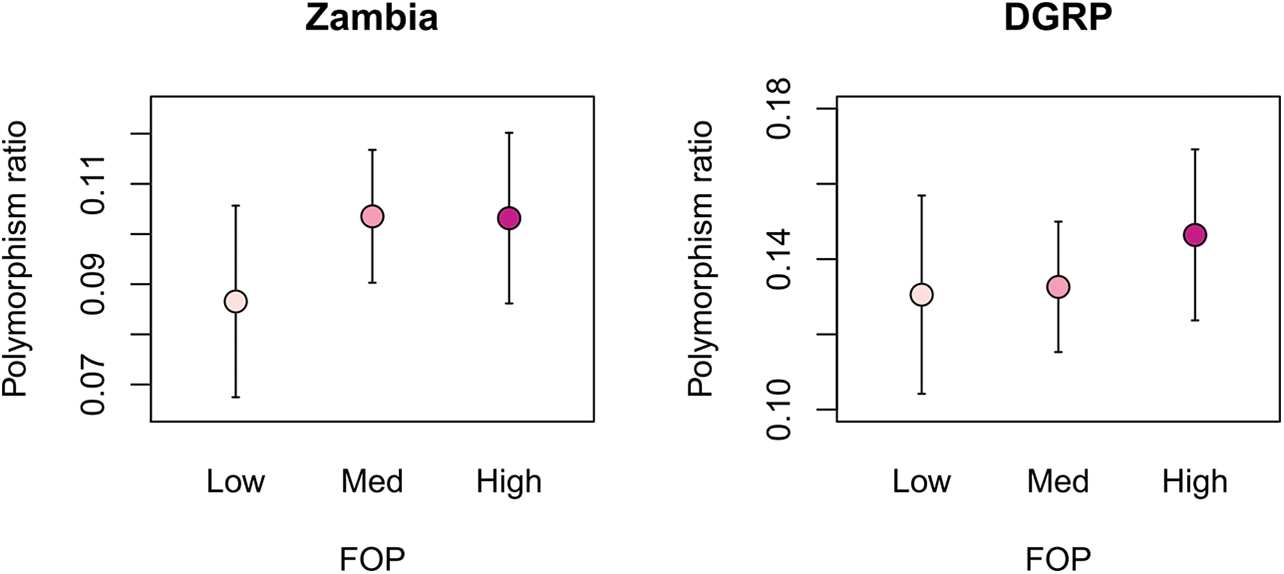
Polymorphism ratio by frequency of preferred codons (FOP). Error bars represent two standard error.

**Figure S5:**
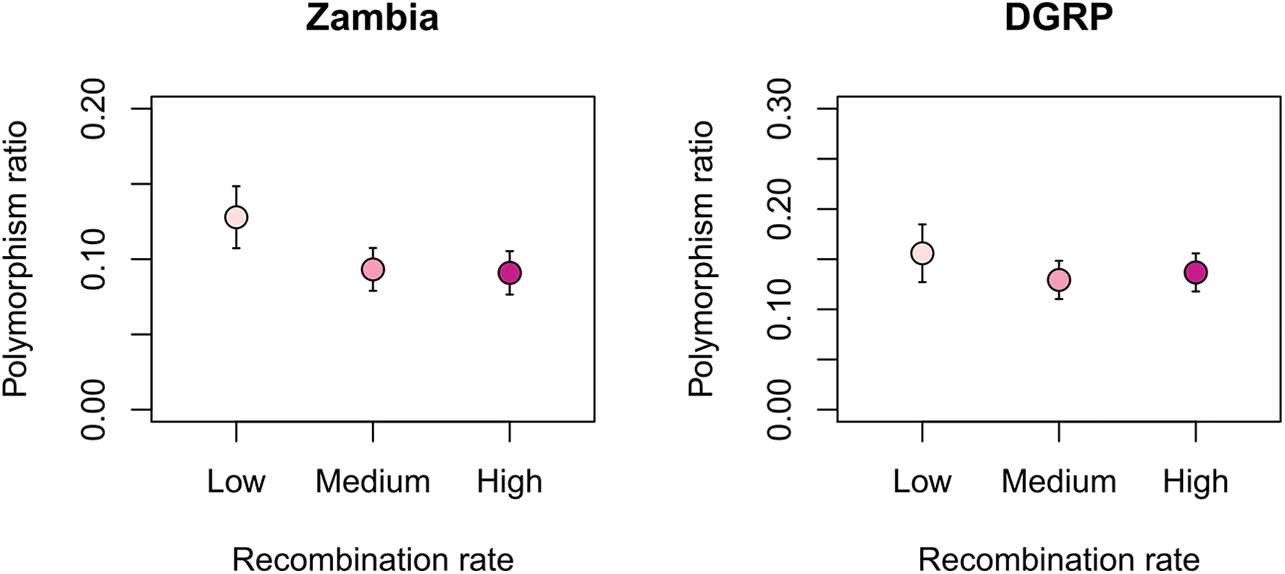
Polymorphism ratio by recombination rate. Error bars represent two standard error.

**Figure S6:**
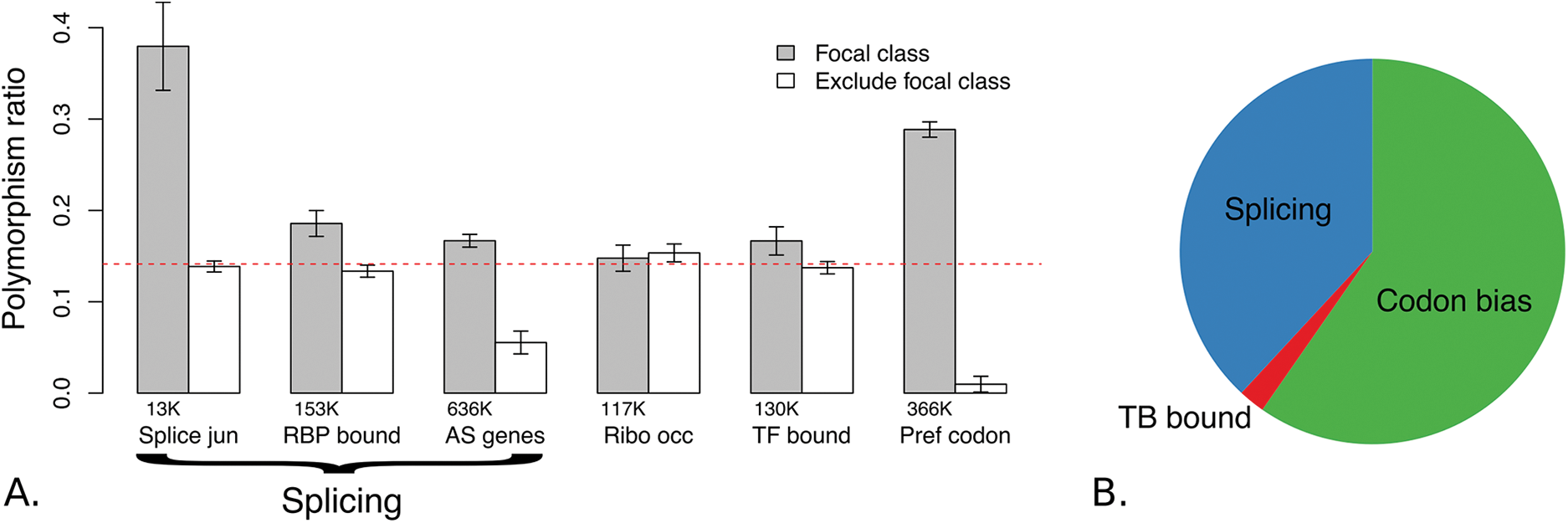
A) Proportion of sites under strong selection as measured by the polymorphism ratio for each class of site (grey) and the dataset excluding the focal class sites (white). The number of sites in a focal class is listed below the corresponding bar. The red dashed line is the polymorphism ratio for the full dataset (DGRP). Error bars represent two standard error. B): Relative proportion of synonymous sites under strong purifying selection due to slicing, codon bias, or transcription factor binding (TFB).

**Table S1:** Six-bin free-alpha model

| Bin number | Frequency bin |
| --- | --- |
| 1 | 1/N |
| 2 | 2/N : 3/N |
| 3 | 4/N : 7/N |
| 4 | 8/N : 15/N |
| 5 | 16/N : (N/2 - 15)/2 |
| 6 | (N/2 - 15)/2 + 1 : N/2 |

## References

Akashi H (1996) Molecular evolution between *Drosophila melanogaster* and *D. simulans*: reduced codon bias, faster rates of amino acid substitution, and larger proteins in *D. melanogaster*. Genetics, 144, 1297–307.

Andolfatto P, Wong KM, Bachtrog D (2011) Effective population size and the efficacy of selection on the X chromosomes of two closely related *Drosophila* species. Genome biology and evolution, 3, 114–128.

Brooks AN, Duff MO, May G, et al. (2015) Regulation of alternative splicing in *Drosophila* by 56 RNA binding proteins. Genome research, 25, 1771–80.

Brown JB, Boley N, Eisman R, et al. (2014) Diversity and dynamics of the *Drosophila* transcriptome. Nature, 512, 393–9.

Bulmer M (1991) The selection-mutation-drift theory of synonymous codon usage. Genetics, 129, 897–907.

Campos JL, Charlesworth B, Haddrill PR (2012) Molecular evolution in nonrecombining regions of the *Drosophila melanogaster* genome. Genome biology and evolution, 4, 278–88.

Campos JL, Zeng K, Parker DJ, Charlesworth B, Haddrill PR (2013) Codon usage bias and effective population sizes on the X chromosome versus the autosomes in *Drosophila melanogaster*. Molecular biology and evolution, 30, 811–23.

Carlini DB (2004) Experimental reduction of codon bias in the *Drosophila* alcohol dehydrogenase gene results in decreased ethanol tolerance of adult flies. Journal of evolutionary biology, 17, 779–85.

Carlini DB, Stephan W (2003) In vivo introduction of unpreferred synonymous codons into the *Drosophila* Adh gene results in reduced levels of ADH protein. Genetics, 163, 239–43.

Chamary JV, Parmley JL, Hurst LD (2006) Hearing silence: non-neutral evolution at synonymous sites in mammals. Nature reviews. Genetics, 7, 98–108.

Chen Sl, Xu My, Hu Sn, Li L (2004) Analysis of immune-relevant genes expressed in red sea bream (*Chrysophrys major*) spleen. Aquaculture, 240, 115 – 130.

Clemente F, Vogl C (2012a) Evidence for complex selection on four-fold degenerate sites in *Drosophila melanogaster*. Journal of evolutionary biology, 25, 2582–95.

Clemente F, Vogl C (2012b) Unconstrained evolution in short introns? - an analysis of genome-wide polymorphism and divergence data from *Drosophila*. Journal of evolutionary biology, 25, 1975–90.

Cooper GM, Stone EA, Asimenos G, Green ED, Batzoglou S, Sidow A (2005) Distribution and intensity of constraint in mammalian genomic sequence. Genome research, 15, 901–13.

Dunn JG, Foo CK, Belletier NG, Gavis ER, Weissman JS (2013) Ribosome profiling reveals pervasive and regulated stop codon readthrough in *Drosophila melanogaster*. eLife, 2, e01179.

Felsenstein J (1974) The evolutionary advantage of recombination. Genetics, 78, 737–56.

Gouy M, Gautier C (1982) Codon usage in bacteria: correlation with gene expressivity. Nucleic Acids Research, 10, 7055–7074.

Grantham R, Gautier C, Gouy M, Jacobzone M, Mercier R (1981) Codon catalog usage is a genome strategy modulated for gene expressivity. Nucleic acids research, 9, r43–74.

Grantham R, Gautier C, Gouy M, Mercier R, Pavé A (1980) Codon catalog usage and the genome hypothesis. Nucleic acids research, 8, r49–r62.

Haddrill PR, Charlesworth B, Halligan DL, Andolfatto P (2005) Patterns of intron sequence evolution in *Drosophila* are dependent upon length and GC content. Genome biology, 6, R67.

Halligan DL, Keightley PD (2006) Ubiquitous selective constraints in the *Drosophila* genome revealed by a genome-wide interspecies comparison. Genome research, 16, 875–84.

Hershberg R, Petrov DA (2008) Selection on codon bias. Annual review of genetics, 42, 287–99.

Hu TT, Eisen MB, Thornton KR, Andolfatto P (2013) A second-generation assembly of the *Drosophila simulans* genome provides new insights into patterns of lineage-specific divergence. Genome research, 23, 89–98.

Ikemura T (1981) Correlation between the abundance of Escherichia coli transfer RNAs and the occurrence of the respective codons in its protein genes: a proposal for a synonymous codon choice that is optimal for the *E. coli* translational system. Journal of molecular biology, 151, 389–409.

Ikemura T (1982) Correlation between the abundance of yeast transfer RNAs and the occurrence of the respective codons in protein genes. Differences in synonymous codon choice patterns of yeast and *Escherichia coli* with reference to the abundance of isoaccepting transfer R. Journal of molecular biology, 158, 573–97.

Jackson BC, Campos JL, Haddrill PR, Charlesworth B, Zeng K (2017) Variation in the intensity of selection on codon bias over time causes contrasting patterns of base composition evolution in *Drosophila*. Genome Biology and Evolution, p. evw291.

Kessler MD, Dean MD (2014) Effective population size does not predict codon usage bias in mammals. Ecology and evolution, 4, 3887–900.

Kliman RM, Hey J (1993) Reduced natural selection associated with low recombination in *Drosophila melanogaster*. Molecular biology and evolution, 10, 1239–58.

Kousathanas A, Keightley PD (2013) A comparison of models to infer the distribution of fitness effects of new mutations. Genetics, 193, 1197–208.

Lack JB, Cardeno CM, Crepeau MW, et al. (2015) The *Drosophila* genome nexus: a population genomic resource of 623 *Drosophila melanogaster* genomes, including 197 from a single ancestral range population. Genetics, 199, 1229–41.

Lawrie D, Petrov Da, Messer PW (2011) Faster than Neutral Evolution of Constrained Sequences: The Complex Interplay of Mutational Biases and Weak Selection. Genome biology and evolution.

Lawrie DS, Messer PW, Hershberg R, Petrov DA (2013) Strong purifying selection at synonymous sites in *D. melanogaster*. PLoS genetics, 9, e1003527.

Li WH (1987) Models of nearly neutral mutations with particular implications for nonrandom usage of synonymous codons. Journal of Molecular Evolution, 24, 337–345.

Mackay TFC, Richards S, Stone EA, et al. (2012) The *Drosophila melanogaster* Genetic Reference Panel. Nature, 482, 173–8.

Marais G, Mouchiroud D, Duret L (2001) Does recombination improve selection on codon usage? Lessons from nematode and fly complete genomes. Proceedings of the National Academy of Sciences of the United States of America, 98, 5688–92.

McVean G, Charlesworth B (1999) A population genetic model for the evolution of synonymous codon usage: patterns and predictions. Genetical Research, 74, 145–158.

Newman ZR, Young JM, Ingolia NT, Barton GM (2016) Differences in codon bias and GC content contribute to the balanced expression of TLR7 and TLR9. Proceedings of the National Academy of Sciences of the United States of America, 113, E1362–71.

Novoa EM, Ribas de Pouplana L (2012) Speeding with control: codon usage, tRNAs, and ribosomes. Trends in genetics: TIG, 28, 574–81.

Parsch J, Novozhilov S, Saminadin-Peter SS, Wong KM, Andolfatto P (2010) On the utility of short intron sequences as a reference for the detection of positive and negative selection in *Drosophila*. Molecular biology and evolution, 27, 1226–34.

Pechmann S, Frydman J (2013) Evolutionary conservation of codon optimality reveals hidden signatures of cotranslational folding. Nature structural & molecular biology, 20, 237–43.

Plotkin JB, Kudla G (2011) Synonymous but not the same: the causes and consequences of codon bias. Nature reviews. Genetics, 12, 32–42.

Post LE, Strycharz GD, Nomura M, Lewis H, Dennis PP (1979) Nucleotide sequence of the ribosomal protein gene cluster adjacent to the gene for RNA polymerase subunit beta in Escherichia coli. Proceedings of the National Academy of Sciences of the United States of America, 76, 1697.

Presnyak V, Alhusaini N, Chen YH, et al. (2015) Codon Optimality Is a Major Determinant of mRNA Stability. Cell, 160, 1111–1124.

Qian W, Yang JR, Pearson NM, Maclean C, Zhang J (2012) Balanced codon usage optimizes eukaryotic translational efficiency. PLoS genetics, 8, e1002603.

Sharp NP, Agrawal AF (2016) Low Genetic Quality Alters Key Dimensions of the Mutational Spectrum. PLoS biology, 14, e1002419.

Sharp PM, Li WH (1986) Codon usage in regulatory genes in *Escherichia coli* does not reflect selection for ‘rare’ codons. Nucleic acids research, 14, 7737–49.

Singh ND, Bauer DuMont VL, Hubisz MJ, Nielsen R, Aquadro CF (2007) Patterns of mutation and selection at synonymous sites in *Drosophila*. Molecular biology and evolution, 24, 2687–97.

Singh ND, Davis JC, Petrov DA (2005) Codon bias and noncoding GC content correlate negatively with recombination rate on the *Drosophila* X chromosome. Journal of molecular evolution, 61, 315–24.

Urrutia AO, Hurst LD (2001) Codon Usage Bias Covaries With Expression Breadth and the Rate of Synonymous Evolution in Humans, but This Is Not Evidence for Selection. Genetics, 159, 1191–1199.

Zeng K, Charlesworth B (2009) Estimating selection intensity on synonymous codon usage in a nonequilibrium population. Genetics, 183, 651–62, 1SI–23SI.

Zeng K, Charlesworth B (2010) Studying Patterns of Recent Evolution at Synonymous Sites and Intronic Sites in *Drosophila melanogaster*. Journal of Molecular Evolution, 70, 116–128.

Zhou J, Liu WJ, Peng SW, Sun XY, Frazer I (1999) Papillomavirus capsid protein expression level depends on the match between codon usage and tRNA availability. Journal of virology, 73, 4972–82.

Zhou Z, Dang Y, Zhou M, et al. (2016) Codon usage is an important determinant of gene expression levels largely through its effects on transcription. Proceedings of the National Academy of Sciences, 113, E6117–E6125.

